# Microbial network inference for longitudinal microbiome studies with LUPINE

**DOI:** 10.1101/2024.05.08.593086

**Authors:** Saritha Kodikara, Kim-Anh Lê Cao

## Abstract

The microbiome is a complex ecosystem of interdependent taxa that has traditionally been studied through cross-sectional studies. However, longitudinal microbiome studies are becoming increasingly popular. These studies enable researchers to infer taxa associations towards the understanding of coexistence, competition, and collaboration between microbes across time. Traditional metrics for association analysis, such as correlation, are limited due to the data characteristics of microbiome data (sparse, compositional, multivariate). Several network inference methods have been proposed, but have been largely unexplored in a longitudinal setting.

We introduce LUPINE (LongitUdinal modelling with Partial least squares regression for NEtwork inference), a novel approach that leverages on conditional independence and low-dimensional data representation. This method is specifically designed to handle scenarios with small sample sizes and small number of time points. LUPINE is the first method of its kind to infer microbial networks across time, while considering information from all past time points and is thus able to capture dynamic microbial interactions that evolve over time. We validate LUPINE and its variant, LUPINE single (for single time point analysis) in simulated data and four case studies, where we highlight LUPINE’s ability to identify relevant taxa in each study context, across different experimental designs (mouse and human studies, with or without interventions, as short or long time courses). We propose different metrics to compare the inferred networks and detect changes in the networks across time, groups or in response to external disturbances.

LUPINE is a simple yet innovative network inference methodology that is suitable for, but not limited to, analysing longitudinal microbiome data. The R code and data are publicly available for readers interested in applying these new methods to their studies.

## 1 Introduction

The microbiome refers to a distinct microbial community that resides within a welldefined habitat, characterized by specific physio-chemical properties (Berg et al, 2020). This ecosystem is composed of billions of bacteria from hundreds to thousands of different taxa (Dicks et al, 2018), forming complex community structures that are challenging to decipher. Given the inherent dynamics of the microbiome, longitudinal studies are essential to gain insights into the impact, variations over time, and responses to external perturbations such as dietary changes or medications on the microbiome. The temporal dimension, while adding complexity, provides richer insights into the dynamic processes, patterns, and interactions within microbial communities, surpassing what cross-sectional studies can offer (Lyu et al, 2023). The advancements in sequencing technologies, coupled with reduced sequencing costs, have enabled researchers to conduct an increasing number of longitudinal microbiome studies.

One objective of microbiome studies is to accurately infer microbial networks from the datasets generated by sequencing technologies. These networks can be viewed as temporal or spatial snapshots of ecosystems (Röttjers and Faust, 2018). In these networks, interactions signify significant associations between taxa, which are typically non-directional in nature (Kurtz et al, 2015). The simplest way to detect associations is via correlation analysis, such as Pearson or Spearman correlation. However, these correlation methods are sub-optimal for microbiome data as they ignore the compositional structure of the data, and lead to spurious results (Gloor et al, 2017). A more valid approach is to use the concept of partial correlation, which effectively measure pairwise associations between variables subject to a constant-sum constraint (i.e. compositional data, Erb, 2020). Existing compositional-aware microbiome network methods have currently focused on single time point studies, rather than longitudinal. Widely used single time point methods are either based on correlation approaches, such as SparCC (Friedman and Alm, 2012) or precision based approaches, such as SpiecEasi (Kurtz et al, 2015). In contrast to correlation-based methods, precision or partial correlation-based methods concentrate on direct associations by removing indirect associations.

Longitudinal microbiome studies enables researchers to create temporal snapshots of microbial networks for a comprehensive view of the microbiome. But network methods for such type of studies are still in their infancy and exhibit diverse analytical objectives (Kodikara et al, 2022). Some methods aim to model relationships between taxa for each individual subject. Other methods aim to model collective microbial interactions across all time points. However, the assumption that microbial interaction stays constant through time, whether on an individual basis or across all subjects, proves limiting. This limitation is particularly evident when there are external interventions, such as alterations in diet or antibiotic usage, which disrupt the stability of these relationships. We have addressed this limitation with a sequential approach, that, to the best of our knowledge, is the first of its kind for longitudinal microbiome studies. We acknowledge the dynamic nature of microbial interactions, especially in the face of interventions, and introduce a flexible framework for the temporal evolution of these interactions.

LUPINE (LongitUdinal modelling with Partial least squares regression for NEtwork inference) combines one-dimensional approximation and partial correlation to measure the linear association between a pair of taxa, accounting for the effects of the other taxa. We developed two variants, one for single time point network inference, and another for longitudinal sequential network inference. The former focuses on inferring the network at a specific time point, in the vein of SpiecEasi (Kurtz et al, 2015) and SparCC (Friedman and Alm, 2012). The latter, however, incorporates information from all previous time points. The inferred network from our method is in the format of a binary graph. In this network, the vertices represent different taxa, and the edges represent significant associations between them. An edge between two taxa indicates that there is a significant connection between them. This connection is visualised as a line between the two nodes in the graph.

In Section 2, we explain the rationale of our approaches along with downstream analyses that can be conducted to compare inferred networks. In the results Section 3, we first benchmark LUPINE and LUPINE single with two state-of-the-art network inference methods (solely designed for single time point microbiome data) on simulated data. Next, through a case study, we demonstrate the robustness of LUPINE compared to LUPINE single. We then showcase the applicability of LUPINE in real data with different experimental designs in four case studies. These experiments varied from human studies to mouse studies, from short to longer time courses, and ranged from cross-sectional to intervention studies. Finally, in the discussion Section 4, we delve into the implications of our findings, special aspects we had to consider when developing LUPINE, and suggest avenues for future research.

## 2 Materials and methods

### 2.1 Method

Our network inference methodology is tailored for the analysis of longitudinal microbiome data, where we assume that the dynamics of taxa interactions are temporal. In order to uncover these intricate relationships, we leverage both past and current information from the data in a sequential manner. Including past information in the model enables insights into inherent variations in microbial interactions across diverse temporal scenarios, including interventions. Our methodology includes three variants that are described in the following sections:

- Single time point modelling: given a pair of taxa, we estimate pairwise partial correlations while accounting for the influence of the other taxa using one-dimensional approximation with principal component analysis (PCA).
- Two time point modelling: given a pair of taxa, we estimate pairwise partial correlations while accounting for the influence of the other taxa with projection to latent structures (PLS) regression (Wold et al, 2001). PLS regression performs dimension reduction such that the covariance between the current and preceding time point data sets is maximised.
- Several time point modelling: given a pair of taxa, we estimate pairwise partial correlations while accounting for the influence of the other taxa using generalised PLS for multiple blocks of data (‘blockPLS’) regression (Tenenhaus and Tenenhaus, 2011; Lê Cao and Welham, 2021). We use blockPLS to maximise the covariance between the current and any past time point datasets.

We assume that individuals within a specific group (e.g. control group) have a common network structure at a particular time point. Therefore, we consider each group separately for studies with multiple sample groups (as illustrated in case studies 1, 3 and 4) to focus on each of their unique microbial characteristics.

#### Notations

In the following, we denote ***X*** an (*n* × *p*) data matrix (where *n* is the number of samples and *p* is the number of taxa); ***X***^*i*^ and ***X***^*j*^ are columns in ***X*** for taxa *i* and *j* ; ***X***^−(*i*,*j*)^ is an (*n* × *p* −2) data matrix excluding taxa *i* and *j* columns in ***X***; ***l*** is a vector of library sizes for *n* samples. In situations where time is of interest (i.e. in longitudinal modelling), all of the above matrices and vectors include a subscript indicating the time point. For example, ***X***_*t*_ represents an (*n* × *p*) data matrix at time point *t*. For simplicity, we will omit the hat notation (i.e. *π* instead of 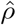) of our estimates. However, note that all estimates approximate some population parameter. Additionally, ∥.∥ denotes the Euclidean norm.

##### Single time point modelling with PCA

We first introduce the single time point approach which we will generalise for multiple time points in the next sections. This type of analysis gives insights of microbial associations at a single time point. It can also be used when there is a large time gap between time points but only one particular time point is of interest.

To assess the strength of the association between two taxa, we use partial correlation – a statistical technique that measures association while controlling for other variables. For example, we estimate the partial correlation between a pair of taxa, denoted by *i* and *j* in Figure 1, to assess the strength of their association while controlling for other taxa. Controlling for these taxa requires regressing taxa *i* and *j* against *p*−2 taxa, which becomes unsolvable when the number of taxa *p* is larger than the number of samples *n* (e.g. in typical microbiome studies *p* = 150−500 and *n <* 50). Hence, the first step in our approach is to calculate a one-dimensional approximation of the control variables (e.g. all taxa except the pair *i* and *j*), through the first principal component.

**Fig. 1.**
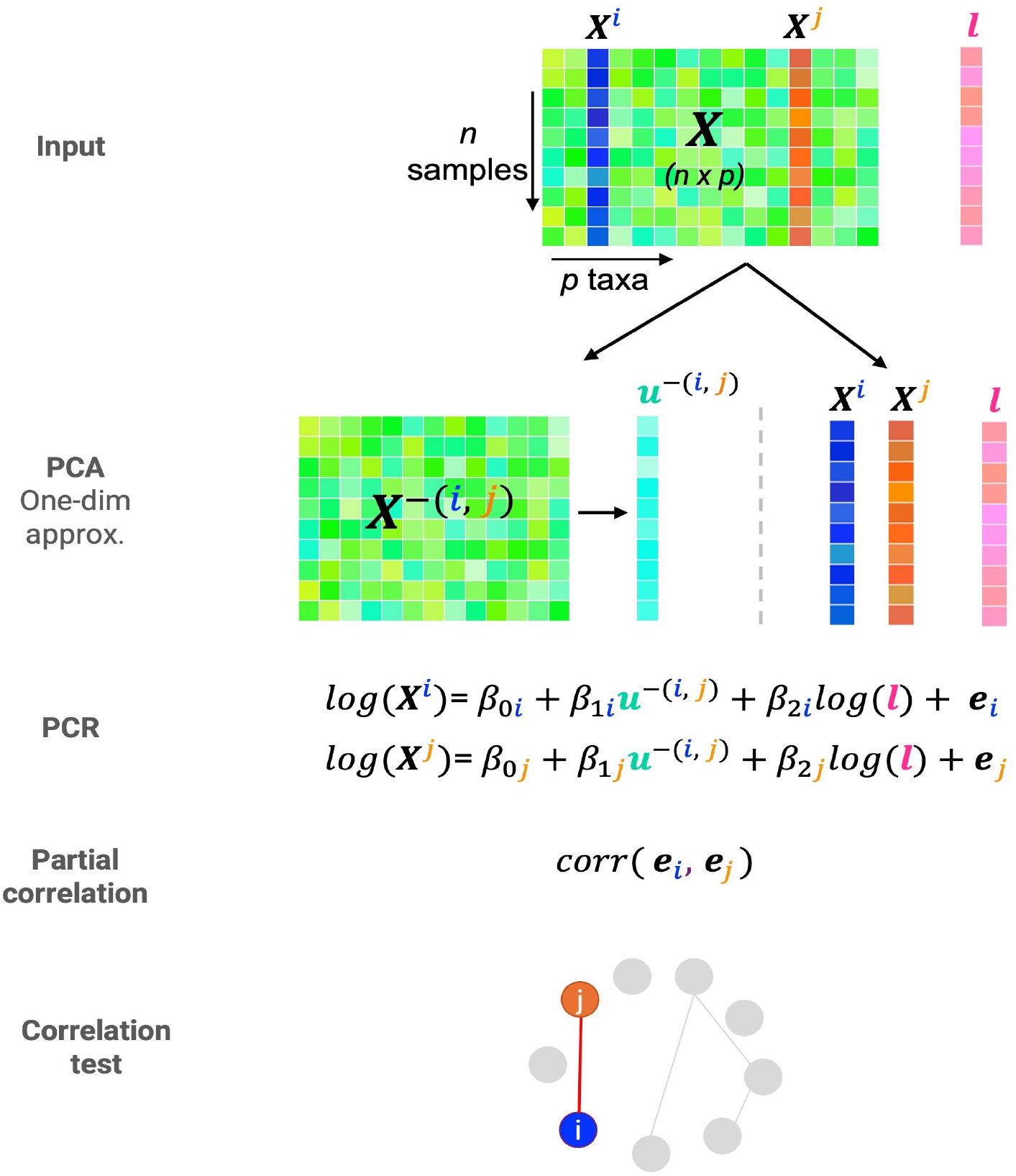
Single time point modelling overview. Consider the (*n* × *p*) input data matrix ***X*** and ***l*** the library size of each sample. First, we use PCA to derive a one-dimensional approximation, ***u***^−(*i*,*j*)^ to control for taxa other than *i* and *j* from the ***X***^−(*i*,*j*)^ data matrix. We then apply two independent log linear regression models (a.k.a. Principal Component Regression) for ***X***^*i*^ and ***X***^*j*^, regressed on component ***u***^−(*i*,*j*)^ and log(***l***) to obtain the residuals ***e***_*i*_ and ***e***_*j*_ . We estimate the partial correlation estimate for taxa *i* and *j* as the correlation between the two residuals. Finally, we test for the statistical significance of the true correlation between ***e***_*i*_ and ***e***_*j*_ to infer whether taxa *i* and *j* are connected in the network. We iterate this process for all pairs of taxa (*i, j*) to obtain a full binary network. Section 2.2 explains how to avoid repeated PCA approximations to improve computational time.

We first start with a generic formulation of PCA (Jolliffe, 2002). The objective function to obtain the first principal component is

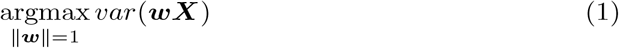

where ***w*** is the *p* dimensional loading vector associated to the first principal component ***u***, with ***u*** = ***wX*** under the constraint that *w* is of unit (norm) 1.

First, we modify Eq (1) to find the first principal component of the control taxa that excludes taxa *i* and *j* as

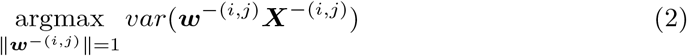

where ***w***^−(*i*,*j*)^ is the (*p* − 2) dimensional loading vector associated to the first principal component ***u***^−(*i*,*j*)^ with ***u***^−(*i*,*j*)^ = ***w***^−(*i*,*j*)^***X***^−(*i*,*j*)^.

Next, we fit two log linear regressions on each of the taxa *i* and *j*. They are regressed against the first principal component, ***u***^−(*i*,*j*)^, similar to principal component regression (PCR, Jolliffe (1982)). PCR combines PCA and multiple linear regression, where the principal components of the explanatory variables serve as predictors rather than using the original variables directly in the regression. Here, we fit two log linear regressions to extract their residuals denoted by ***e***_*i*_ and ***e***_*j*_:

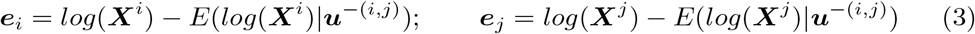

where *E*(.| ***u***^−(*i*,*j*)^) is the conditional expectation with respect to the first principal component ***u***^−(*i*,*j*)^. We use a log linear regression as the response variables are counts. This is similar to the bias-corrected version of ANCOM (Lin and Peddada, 2020) that uses a linear regression framework based on log-transformed taxa counts. We add an offset of 1 to zero counts to ensure that the log transformation is valid. Additionally, in Supplementary Section S4.2, we assess the impact of using log-transformed counts in Eq 2.

In addition, to correct for library size (i.e. the variation in the total sequence reads), we modify Eq 3 and include an offset, similar to the approach of Zhang et al (2018, 2020):

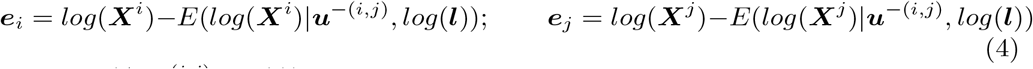

where *E*(.|***u***^−(*i*,*j*)^, *log*(***l***)) is the conditional expectation with respect to the first principal component ***u***^−(*i*,*j*)^ and the log library size log(***l***).

We then quantify the strength of association between taxa *i* and *j* based on partial correlation using the residuals ***e***_*i*_ and ***e***_*j*_. Partial correlations reveal direct (i.e. conditionally independent) associations between two variables while controlling for the other variables. We calculate the partial correlation between the pair of taxa *i* and *j* as

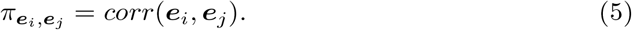

Finally, we test whether the true correlation 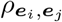,between ***e***_*i*_ and ***e***_*j*_ (estimated with 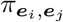) is significantly different from zero. The correlation test assumes that when the correlation is zero, the test statistic 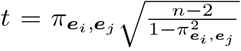,follows a normal distribution. Thus, we test the hypothesis 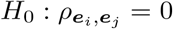 versus 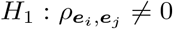 using a t-distribution with *n*−2 degrees of freedom. We then consider that a pair of taxa is connected in a network if the associated p-value from the correlation test is less than a chosen significance level (e.g. 0.05). We apply this approach to all possible pairs of taxa to obtain a binary network summarising the pairwise connections between all taxa. In Section S4.1, we compare this analytical approach with a permutation-based p-value calculation.

To quantify the uncertainty of the correlation test p-value, we use bootstrapping, which helps determine the significance of a correlation. We resample the data set 1,000 times and calculate the p-values from correlation tests for each iteration. This process generates a confidence interval for each p-value, allowing us to better assess the significance of the correlation. We demonstrate this approach in case study 4.

In the pseudo-algorithm 1 and Figure 1, we outline the different steps in LUPINE single. In Section 2.2, we explain how we improve computational time by avoiding repeated PCA approximations.

###### Algorithm 1

Single time point scheme

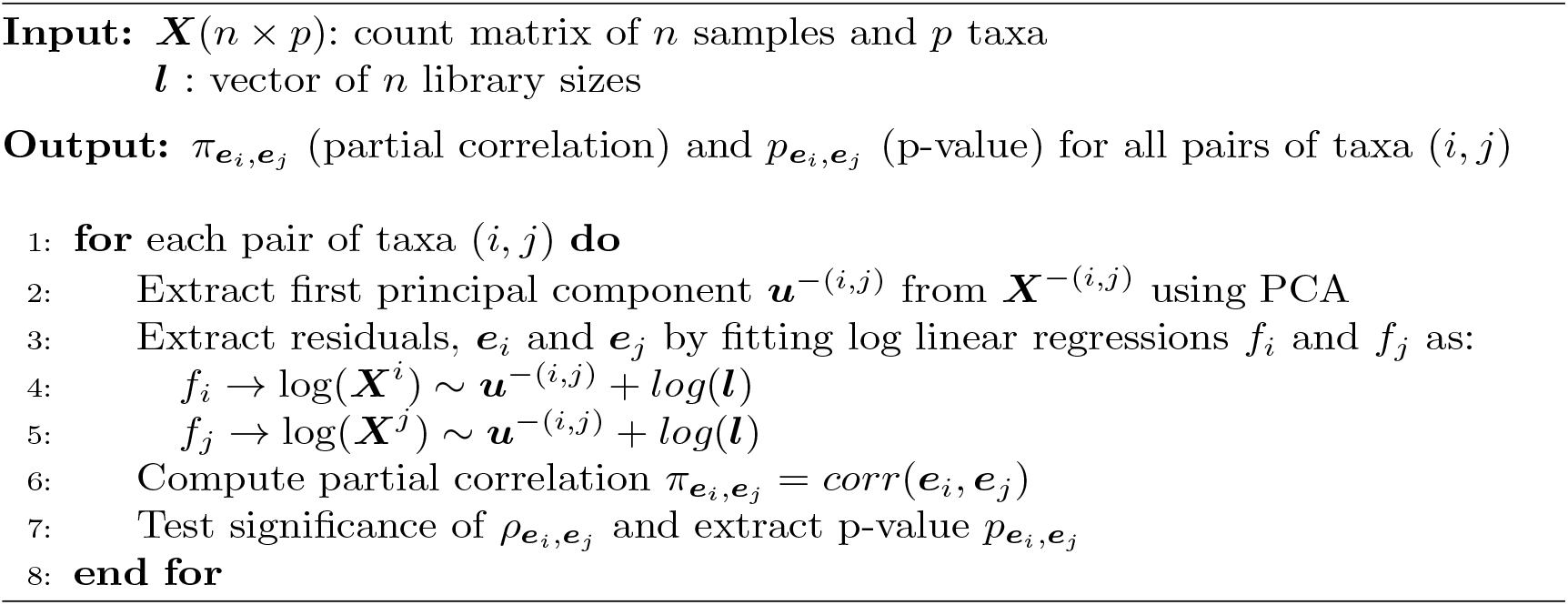

##### Two time point modelling with PLS regression

Here we use PLS to obtain a one-dimensional approximation between two datasets from two time points (Figure 2). PLS is a projection-based method that maximises the covariance between the latent components associated to two data sets while managing correlated information (Wold et al, 2001). The PLS latent variables (or components), are linear combinations of variables from each data set. These latent variables aim to uncover subtle biological effects not evident otherwise if each data set is considered independently (Lê Cao and Welham, 2021).

**Fig. 2.**
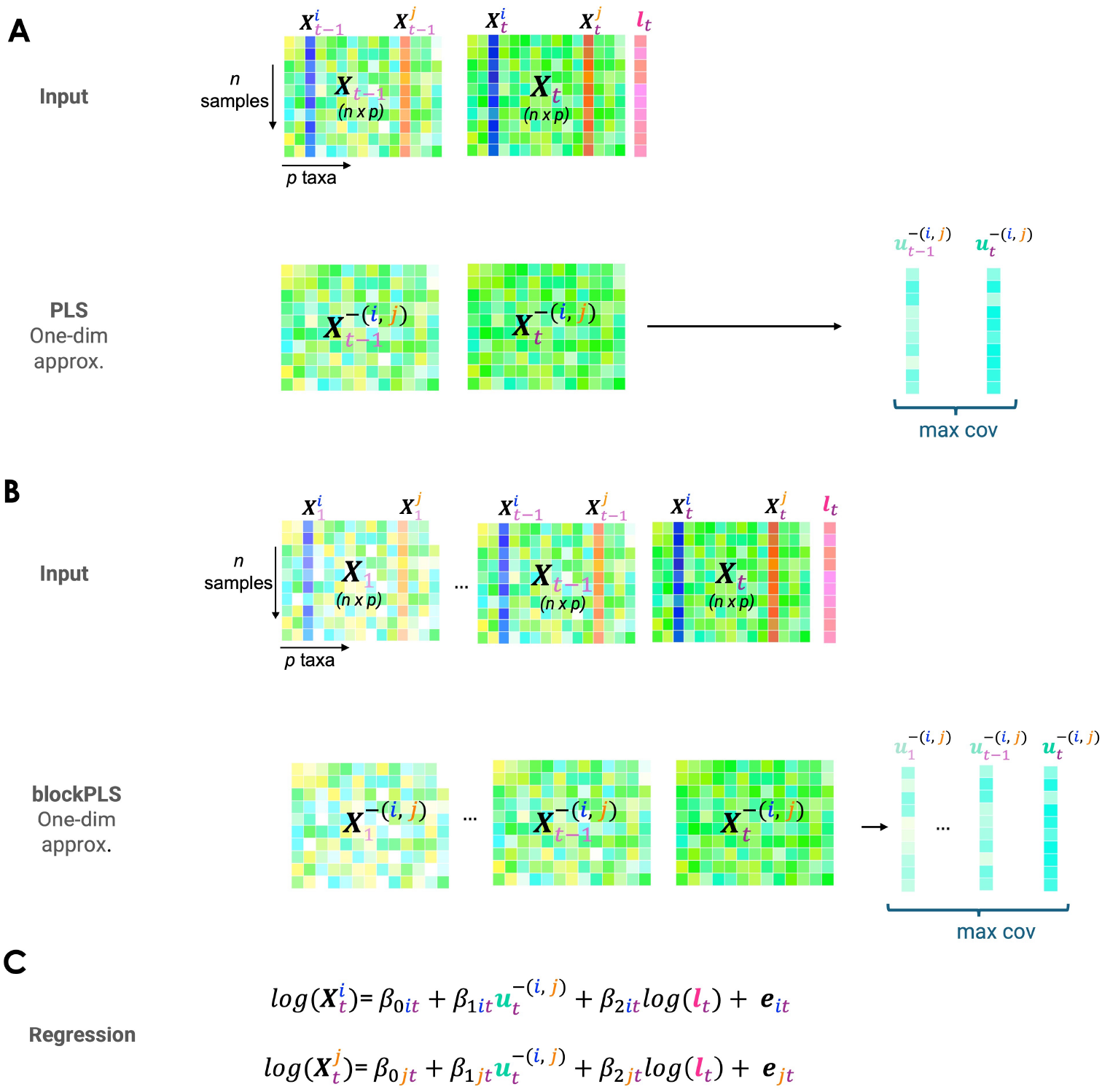
Two or more time points modelling overview. Consider as input the (*n* × *p*) data matrices for **A** two time points or **B** more than two time points (denoted ***X***_1_ …, **X**_*t*_) and their respective library size ***l***_*t*_ at time *t*. Similar to single time point modelling presented in Figure 1, first, we derive a one-dimensional approximation, 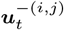 to control for taxa other than *i* and *j*. Instead of using PCA as previously for a single time point, we use either **A** PLS maximising the covariance with the control taxa at the preceding time *t* − 1, or **B** blockPLS maximising the covariance with the control taxa at all previous times 1, …, *t* − 1. **C** We then fit two independent log linear models for 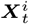 and 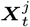, regressed on 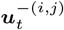 and log(***l***_*t*_) to compute the residuals ***e***_*it*_ and ***e***_*jt*_ following either **A** or **B**. The remainder of the modeling process, including the partial correlation calculation and the correlation test, is identical to that in Figure 1.

The objective function for the first dimension of PLS is

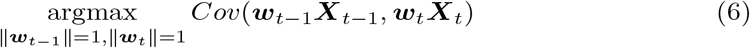

where ***w***_*t*−1_ and ***w***_*t*_ are the two *p* dimensional loading vectors associated to the first latent components ***u***_*t*−1_ = ***w***_*t*−1_***X***_*t*−1_ and ***u***_*t*_ = ***w***_*t*_***X***_*t*_.

We modify Eq 6 to find the first latent components of the controlling taxa at time point *t* that has a maximum covariance with the preceding time point latent component as

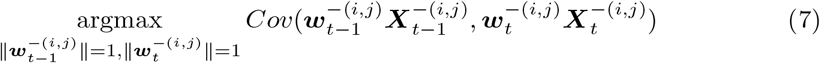

where 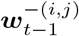 and 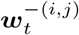 are the two (*p* − 2) loading vectors associated to the first latent components 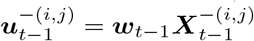 and 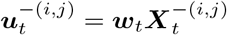.

To infer the network at time point *t* using the information from the preceding time point (*t* − 1), we fit two log linear regressions. These regressions are fitted on 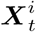 and 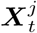 that are regressed against the first latent component 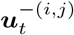, and the log library size log(***l***_*t*_). Note that 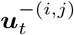 differs from the single time point approach 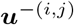 as we maximise the covariance with the preceding time point latent component 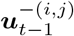 . The updated Eqs 4 and 5 become

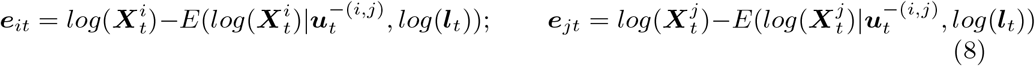

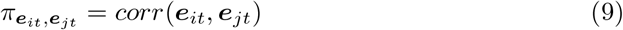

where 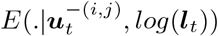 is the conditional expectation with respect to the first latent component 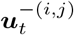 and the log library size, log(***l***_*t*_) at time point *t*; and ***e***_*it*_ and ***e***_*jt*_ are the residuals from the two log linear models.

Similar to the single time point scheme described above, we test for the strength of association between taxa *i* and *j* with a correlation test, and obtain a binary network summarising the pairwise connections between all taxa after controlling for all other taxa from current time point and previous time point.

##### Multiple time point modelling with blockPLS regression

We use blockPLS, an extension of PLS, to calculate a one-dimensional approximation that maximises the covariance between latent components associated to the current time point, and all past time points. BlockPLS is based on generalised PLS (Tenenhaus and Tenenhaus, 2011; Lê Cao and Welham, 2021) that involves regressing the response matrix (e.g. ***X***_*t*_) on multiple datasets (***X***_1_, ***X***_2_, …, ***X***_*t*−1_).

The objective function for the first dimension of blockPLS is

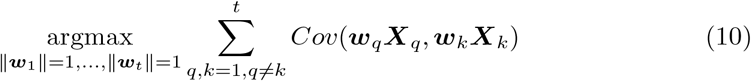

where ***w***_*q*_ is a *p* dimensional loading vector associated to the first latent component ***u***_*q*_, where ***u***_*q*_ = ***w***_*q*_***X***_*q*_.

We modify Eq 10 to find the first latent components of all controlling taxa at time point *t* that has a maximum covariance with previous time points latent components as

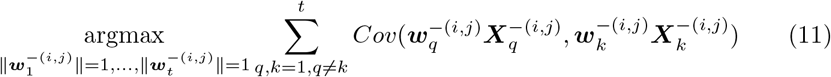

where 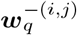 is a *p* − 2 dimensional loading vector associated to the first latent component 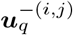 at time point *q*, with 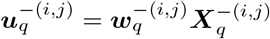.

To infer a network at any time point *t* using all time points until time point *t*, we fit two log linear regressions on the counts of 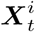 and 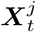 against the first latent component at 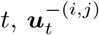, and log library size at *t*, log(***l***_*t*_). However, we need to keep in mind that the latent component 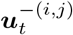 maximises the covariance with all previous time points latent variables, 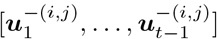, thus effectively taking into account all past time points. We then use Eqs 8 and 9 to quantify and test the statistical significance of the partial correlation between taxa *i* and *j*.

Pseudo-algorithm 2 outlines the different steps in either two time points and multiple time points modelling.

###### Algorithm 2

Multi time point scheme

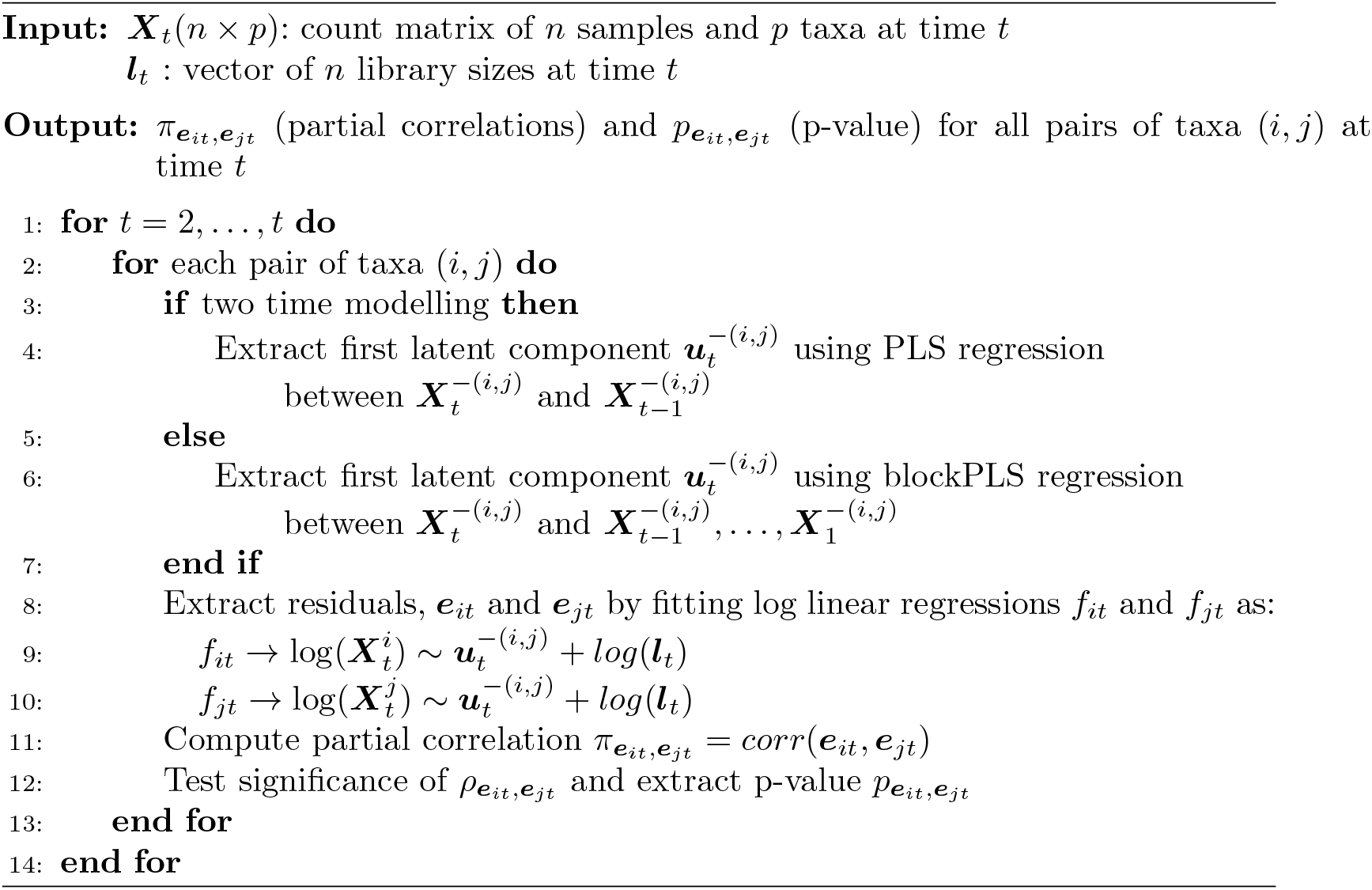

Note that in both PCA and PLS versions we centre and scale each taxon to avoid scaling issues.

### 2.2 Improvement in computational time

Our approach requires to compute one-dimensional approximation for each pair of taxa (*i, j*), i.e. *p*∗ (*p* −1)*/*2 times. Using matrix decomposition principles, we propose instead to approximate the full data matrix ***X*** with the first one-dimension component ***u*** that summarises most of the variation in ***X***, and its associated loading vector that are obtained from PCA, PLS, or blockPLS:

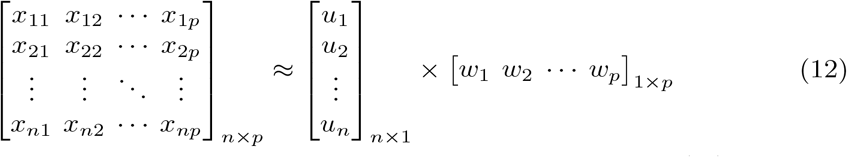

Using the approximation from Eq 12, we then can calculate the ***u***^−(*i*,*j*)^ vector by setting the loading coefficients ***w***_*i*_ and ***w***_*j*_ of the pair of taxa (*i, j*) to zero:

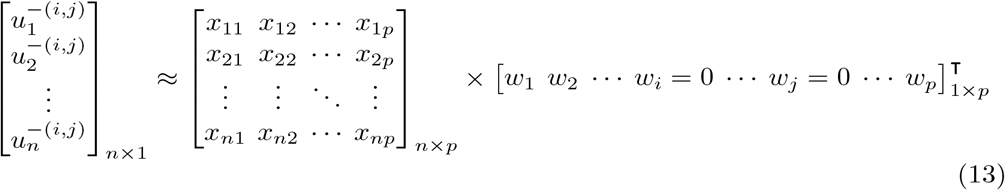

This allows us to obtain the vectors ***u*** and ***w*** only once. We show the efficacy of this approximation in Supplementary section S1.

### 2.3 Comparisons to other network inference methods

Existing longitudinal network methods either infer networks for a single individual or derive a single network from all data (Kodikara et al, 2022), which can be problematic when we expect changes in taxa associations over time. LUPINE infers networks sequentially to include the information learned across time. We compared our results with two state-of-the-art single time point network inference methods, SpiecEasi (Matchado et al, 2021) and SparCC (Matchado et al, 2021), using simulations where the true network is known. SpiecEasi and SparCC represent two distinct branches of network analysis. Specifically, SparCC employs a correlation-based approach, while SpiecEasi uses a conditional dependence approach. The SpiecEasi method further includes two methods based on different graphical model estimators, representing different types of network analysis: (i) neighborhood selection and (ii) sparse inverse covariance selection, which we will refer to as ‘SpiecEasi mb’ and ‘SpiecEasi glasso’, respectively.

### 2.4 Measures for network comparisons

We used two distinct measures to identify similarities and dissimilarities among the inferred networks: a network distance measure for quantitatively evaluating the network topology (see Figure 3B1 - B2), and a node-wise measure for assessing the influential nodes in the network (see Figure 3C1 - C2). Additionally, we measured the differences between network pairs using hypothesis testing (Figure 3D).

**Fig. 3.**
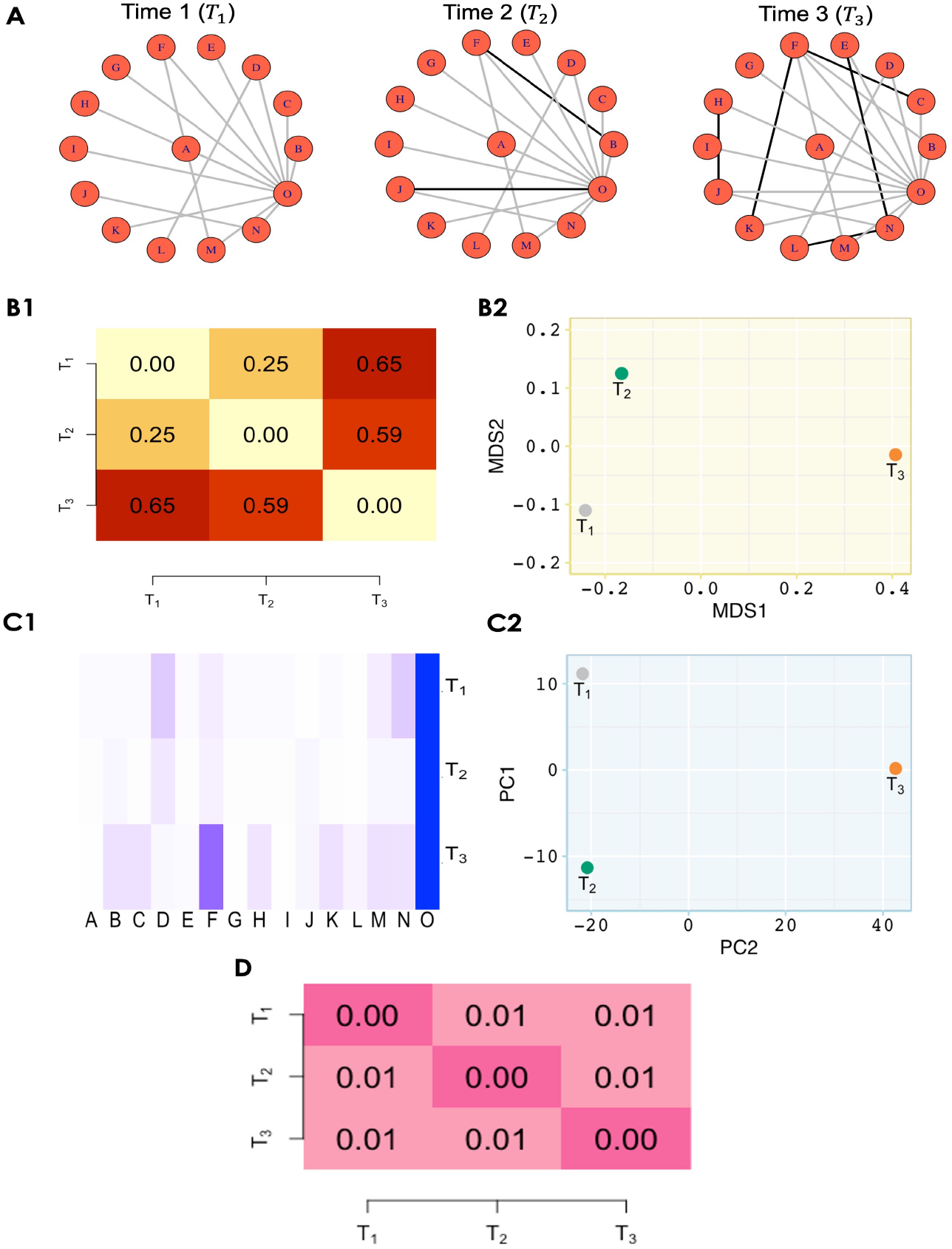
Toy example for network comparison. **A** Consider three networks across three time points, where nodes represent taxa and edges represent significant association between taxa. Black edges denote new connections compared to the preceding time point. **B** Network dissimilarity through graph diffusion distance (GDD). **B1** Pairwise GDD distances reveal that the network at *T*_3_ is dissimilar to the networks at *T*_1_ and *T*_2_. **B2** Transformation of pairwise GDD into a lower-dimensional space using classical Multidimensional Scaling (MDS) represents the separation of time 3 network from the other two networks. **C** Network discrimination through influential nodes (i.e taxa). Node influence is computed by integrating local, semi-local, and global measures. **C1** Heatmap showing the node influence (more influential = darker blue shades). For example, node ‘O’ is influential in all time points, but node ‘F’ is more influential at *T*_3_ rather than *T*_1_ and *T*_2_. **C2** Principal Component Analysis (PCA) plot to represent the networks based on their respective node influence. Similar to what we observed with the GDD distance, the largest source of variation is explained between the network at *T*_3_ and the networks at earlier time points. **D** A heatmap of p-values from the correlation tests between each pair of networks indicates that all three networks are correlated.

#### Pairwise distance between networks

We use Graph diffusion distance (GDD) (Hammond et al, 2013) to measure pairwise differences in network typologies. GDD assesses the average similarity between two networks by exploring the information flow and connectivity within their structures, based on the heat diffusion process on graphs. The GDD score is calculated by searching for a diffusion time that maximises the Frobenius norm of the difference between the diffusion kernels. By calculating the GDD for all pairs, we obtain a pairwise distance matrix representing the network distances between pairs of graphs. We then transform these pairwise distances into a lower-dimensional space using classical Multidimensional Scaling (MDS) (Torgerson, 1958). MDS maintains pairwise dissimilarities between objects. As a result, in the MDS plot, points that represent each network are deemed similar if they are positioned closely, and dissimilar if they are spaced further apart. This visual technique allows us to compare multiple networks at once (Figure 3B2).

#### Node influence

To assess influential nodes in the network, we use the Integrated Value of Influence (IVI) algorithm that integrates local (degree centrality and ClusterRank), semi-local (neighbourhood connectivity and local H-index), and global (betweenness centrality and collective influence) measures (Salavaty et al, 2020). IVI combines these measures to capture both local prominence as well as the broader impact of the nodes. After calculating the IVI values for all nodes, we perform a PCA on the resulting matrix, where the rows represent different networks, and the columns represent different nodes (here taxa). While MDS aims to preserve pairwise distances or dissimilarities as much as possible, PCA prioritise explaining the maximum variation in the data and highlighting strong patterns (Figure 3C2).

#### Correlation test between two networks

To evaluate correlations between two network adjacency matrices, we first calculate pairwise Hamming distances for each matrix (Waggener and Waggener, 1995). This step allows us to compare differences in connections between nodes. Next, we apply the Mantel test (Mantel, 1967), a permutation-based method, to assess whether the correlation between these distance matrices is statistically significant. By comparing the observed correlation to a distribution generated from random permutations, the Mantel test evaluates its significance, providing a robust measure of the relationship between the two matrices.

### 2.5 Simulation and case studies

In this section, we outline our simulation strategy and detail the four case studies analysed with LUPINE.

#### 2.5.1 Simulation study

In our simulation, we used two realistic networks based on our high-fat high-sugar (HFHS) case study described in Section 2.5.2. The two networks were inferred using the SpiecEasi method (Kurtz et al, 2015) where concatenated the data across days for each diet. Due to computational constraints, we restricted our simulation to the subnetwork of the *Bacteroidales* order, resulting in 54 nodes. The networks used across the two time periods are represented in Figure 4. We simulated data using the network in Figure 4A until day 5, then transitioned to the network in Figure 4B from day 6 to day 10 to reflect a network change. We used a multivariate Poisson distribution to simulate the count data. For each simulation, a new correlation matrix was generated from the two networks. To explore the impact of sample size, we varied the sample size from 23 to 50, and then to 120. For each sample size, 50 data sets were simulated, each with 54 taxa and 10 time points. More details on the simulation strategy are provided in the Supplemental Section S2.

**Fig. 4.**
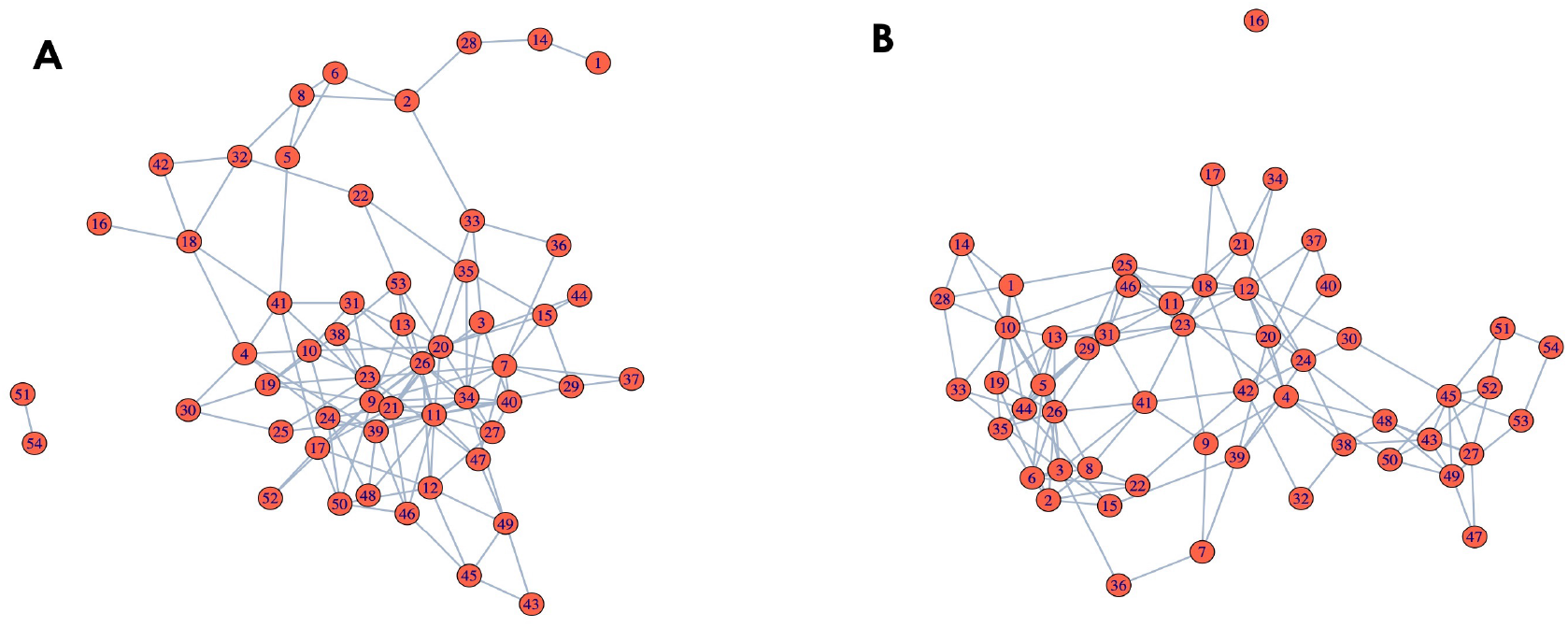
Networks used in simulation. Inferred network from SpiecEasi for **A** normal diet and **B** HFHS diet among *Bacteroidales*. These networks form the basis of our simulation for the two time periods (i.e. from day 1 to 5 and from day 6 to 10).

Using simulated data, we also conducted a sensitivity analysis in Supplemental Section S4 to evaluate the robustness of LUPINE and LUPINE single under various conditions. The analysis was designed to determine whether the model’s performance remains consistent when different parameters or assumptions are altered.

#### 2.5.2 Case studies

We analysed three 16S rRNA amplicon datasets and one metagenomic dataset, the experimental designs of which are outlined in Table 1. For the 16S datasets which were available as raw counts, we grouped data by ‘group’ and ‘time point’, and calculated the relative abundance of each taxa within these specific groups and time points. Any taxa with a relative abundance less than 0.1% across all groups and all time points were excluded. This filtering process, while significantly reducing the number of taxa, aligns with the goal of conducting a more focused analysis (Sankaran and Jeganathan, 2023). However, we only calculated the partial correlations for taxa that had a mean relative abundance exceeding 0.1% at a given time point and group. This approach helps us avoid identifying connections between taxa with low abundance at certain time points in specific groups. As a result, there were no edges in the network from the taxa with low abundance at that time point and group. When we fit a longitudinal model on a given time point, we only consider the taxa filtered on that specific time point and group, and across the past time points.

**Table 1.** Summary of Experimental Frameworks: Key Parameters for Each Case Study. Number of taxa are indicated after the filtering step described in Section 2.5.2.

| Name | Sample size | Number of taxa | Time points | Collection | Interventions |
| --- | --- | --- | --- | --- | --- |
| High-fat high-Sugar (HFHS) | HFHS = 23<br>Normal = 24 | 212 | 4 | Days<br>(0, 1, 4, 7) | NA |
| Vancomycin-resistant Enterococcus faecium (VREfm) colonisation | 9 | 126 | 11 | Days<br>(0-2, 5-9, 12-14) | Ceftriaxone antibiotic treatment (days 6-7)<br>VREfm colonisation (day 9) |
| Diet study | Plant diet = 10<br>Animal diet = 10 | 311 | 15 | Days<br>(-4, -3, -2, -1, 0, 1, 2, 3, 4, 5, 6, 7, 8, 9, 10) | Diet intervention (days 0-4) |
| Pre-diabetic study | MDE diet <sup>a</sup> = 96<br>PPT diet <sup>b</sup> = 93 | 91 | 2 | Months<br>(0,6) | Diet intervention (months 0-6) |

<sup>a</sup> mediterranean diet, <sup>b</sup> personalized postprandial glucose-targeting diet

##### Case Study 1: HFHS (a four time point case-control mouse study)

We investigated the effect of a high-fat high-sugar (HFHS) diet on the mouse microbiome, as outlined in Susin et al (2020). To assess the diet effect on the gut microbiome, 47 C57/B6 female black mice were fed with either an HFHS or a normal diet. Fecal samples were collected at Days 0, 1, 4 and 7.

The raw data contained 1,172 taxa across all four time points. We removed one outlier mouse from the normal diet group with a very large library size. After filtering, we modelled 102,107,105, and 91 taxa respectively for the normal diet for each day, and 99, 147, 85, and 92 taxa for the HFHS diet.

##### Case Study 2: VREfm (an eleven time point mouse study with two interventions)

Mu et al (2020) investigated the functional roles of the gut microbiome during Vancomycin-resistant Enterococcus faecium (VREfm) colonisation. To assess the bacterial community composition in the murine gut, 9 C57BL/6 mice (co-housed wild-type males) were monitored. Fecal samples were collected over 14 days from the same three mice. Over the course of 14 days, the mice underwent two interventions: (1) administration of ceftriaxone treatment at a concentration of 0.5g/liter in the drinking water over 2 days on days 6 and 7, and (2) colonisation with a dosage of (1 × 106) VREfm ST796 on day 9. The days before the antibiotic administration were considered as the naive phase of the experiment. The two days when the antibiotic was given were considered the antibiotic phase, and all other days after that were considered the VRE phase.

The data contained 3,574 taxa across eleven time points. Day 8 was excluded from the analysis due to insufficient taxonomic counts for two mice. After filtering, we considered the following number of taxa in each phase: naive (53, 63, 58, 72), antibiotic (34, 22) and VRE (10, 15, 15, 15).

##### Case Study 3: Diet (a fifteen time point case-control human study with one intervention)

David et al (2014) investigated the human gut microbiome responses to a short-term diet intervention. To evaluate the dynamics of the diet intervention, they assigned 10 participants to each diet intervention, namely the ‘plant-based’ and ‘animal-based’ diets for five consecutive days. The study participants were observed for four days before the diet intervention, a period referred to as the ‘baseline period’ to study their regular eating habits. After intervention, a six-day ‘washout period’ was also observed to assess microbial recovery. Data were obtained from the MITRE repository (Bogart et al, 2019). The data contained 17,310 taxa across all time points. As taxonomic details were lacking, we used the sequence information along with dada2 reference database “silva nr99 v138.1 wSpecies train set.fa.gz” for taxonomic classification (Callahan et al, 2016).

After filtering, we retained the following number of taxa in the plant-based diet group for each period: baseline (105, 100, 111, 103), intervention (118, 116, 116, 114, 114), and washout (121, 120, 90, 96, 115, 109); and in the animal-based diet group: baseline (120, 109, 128, 115), intervention (113, 113, 108, 105, 110), and washout (112, 104, 106, 105, 107, 123). Across the duration of the experiment, only 8-15 samples were obtained from each participant because of unsuccessful sampling at certain time points. Thus, we imputed the missing values using a cubic splines on the centred log ratio (clr) transformed data. This was motivated by the linear interpolation and cubic splines interpolation adopted in Sankaran and Jeganathan (2023); Ruiz-Perez et al (2021). Since, clr transformation accounts for library size differences, we used Eq 3 instead of Eq 4 in our modelling.

##### Case Study 4: Pre-diabetic (large human case control study)

Shoer et al (2023) investigated the impact of a personalized postprandial glucose-targeting diet (PPT) and Mediterranean diet (MED) on gut microbiome on 200 pre-diabetic individuals. MED is the standard of care for pre-diabetes. Participants provided fecal samples before and after a six month diet intervention.

The data contained 378 fecal gut samples with 605 species-level genome bins (SGBs) given as unique relative abundance (URA, Rothschild et al, 2022). After removing *Archaea* and unknown kingdoms, we were left with 505 features, but with many missing values. We only kept features with less than 25% missing values across all samples, resulting in 91 SGBs.

## 3 Results

We first evaluated our two approaches against two widely recognised network inference methods (described in Section 2.3) for cross sectional microbiome data. Using simulated data, our objective was to demonstrate the effectiveness and computational efficiency of our proposed approaches compared to commonly used single time point methods. We then highlighted the advantages of LUPINE over its single time point counterpart LUPINE single in the HFHS study.

After this benchmark step, we applied LUPINE in four case studies. In the longitudinal case-control mouse study, we evaluated whether the difference in diets could be detected in our network models. In the mouse study with two interventions, we evaluated the effectiveness of LUPINE in detecting abrupt changes. In the human casecontrol study, we assessed the ability of LUPINE to handle inherent high variability typical of human studies across fifteen time points. Finally, in the metagenomic human case-control study, we assessed LUPINE’s capacity to generate biological inferences in a large human study with two time points.

### Benchmarking analysis

#### Simulation study: LUPINE methods outperformed existing single time point methods

We first compared our proposed longitudinal method (LUPINE) and its single time point version (LUPINE single), with two SpiecEasi models and SparCC. Note that all methods except LUPINE were designed for single time point modeling.

We assessed model performance across 50 simulated datasets, focusing on scenarios where the number of samples was smaller than the number of variables. We used two metrics: the receiver operating characteristic (AUC-ROC) curve and the precision-recall characteristic (AUC-PRC) curve. True positives were defined as the edges present in the network for each time point. For LUPINE single, LUPINE, and SparCC, we used p-values from the correlation test to infer edges between pairs of nodes. A higher value for 1 −p-value indicated a greater likelihood of an inferred edge, with the value reflecting the strength of the connection between a pair of taxa. For the two SpiecEasi methods, stability was used to measure the strength of the inferred connection.

The AUC-ROC curve captures the trade-off between sensitivity (true positive rate) and specificity (false positive rate) across different thresholds (1−p-value or stability), providing a broad overview of model performance. In contrast, the AUC-PRC curve focuses on precision (positive predictive value) versus recall (true positive rate), which is particularly informative for scenarios where the number of true edges is much smaller than the number of non-edges. By analysing these two metrics, we gained insights into the model’s ability to distinguish true from spurious edges depending on network sparsity. Higher values in both metrics signified better model performance, with AUC-ROC evaluating overall discriminatory power and AUC-PRC evaluating precision and recall characteristics.

Our proposed methods and SparCC outperformed SpiecEasi with higher AUC-ROC values (Figure 5A). However, all methods had similar AUC-PRC values (Figure 5B). Additionally, both our approaches were computationally more efficient than SparCC (Figure 5C). When the sample size increased, LUPINE and LUPINE single led to superior model performance based on AUC-ROC and AUC-PRC values (Figures S3.1 and S3.2).

**Fig. 5.**
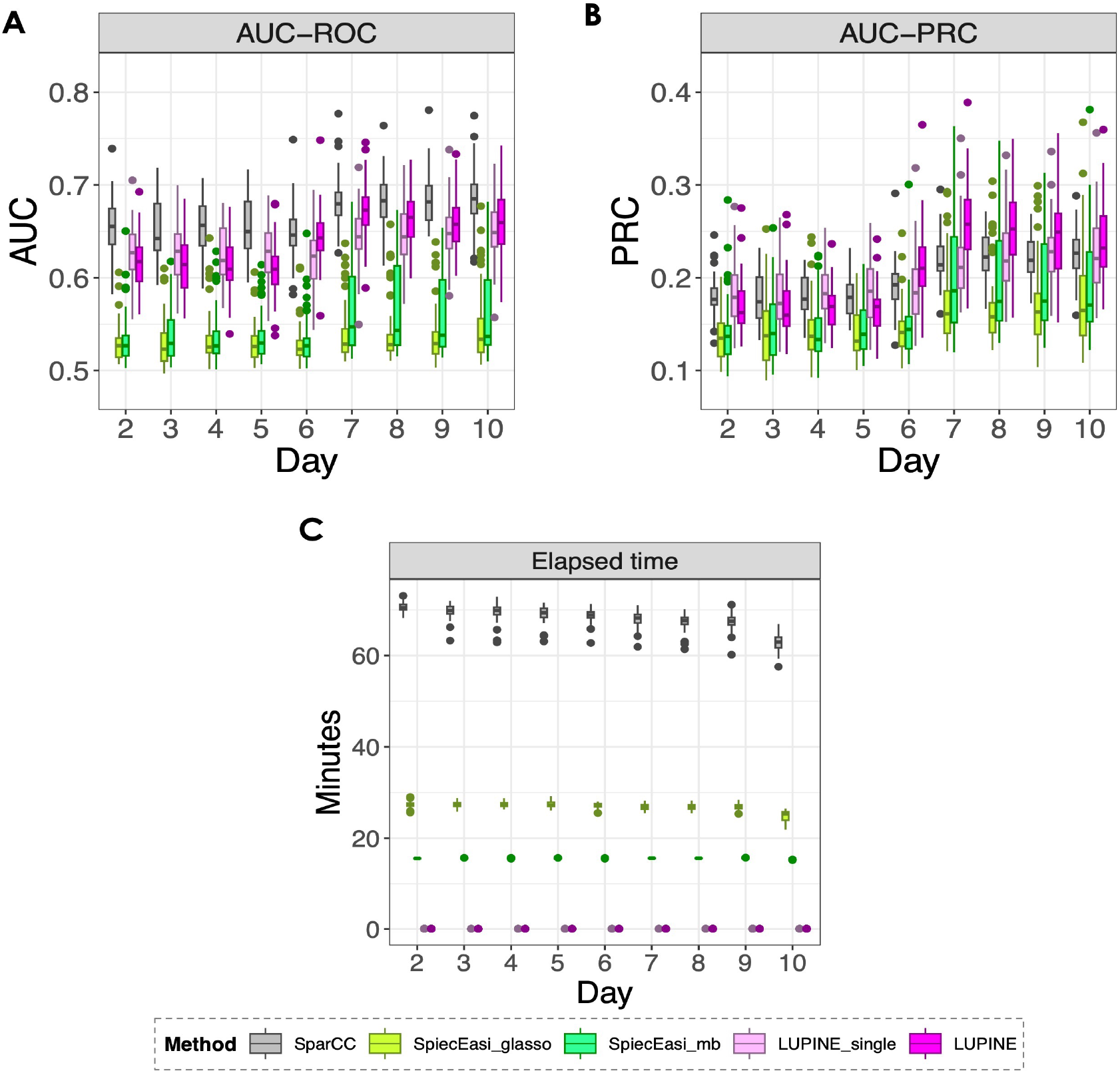
Simulation results (n = 23). Box plots of the Area Under the Receiver **A** Operating Characteristic (AUC-ROC) **B** Precision-Recall Curve (AUC-ROC) over time for different methods. Although all methods performed similarly in terms of AUC-PRC values, both LUPINE models and SparCC outperformed the two SpiecEasi methods based on AUC-ROC values. **C** Elapsed time for each method, showing the superior computational performance of the two LUPINE methods.

Next, we compared the inferred networks across time points, as explained in Section 2.4. For SparCC and our methods, taxa connections were determined via a p-value cutoff (this was not required for SpiecEasi methods, as they inherently output a binary network).

To compare networks based on structure, we computed the pairwise GDDs for all inferred networks across time points, then averaged these distances for each time point across all simulations. In our simulation (described in Section 2.5.1), we generated two stable networks that were distinct, across two time periods (days 2-5 and 6-10). Thus, we expect the inferred networks to be similar within each period, and dissimilar between the two periods. However, SpiecEasi displayed the largest variation between the inferred and the actual networks in the MDS plot of the pairwise GDDs (Figure 6A2 - A3). In contrast, our two proposed methods (Figure 6A4 - A5) and SparCC (Figure 6A1) showed the largest between-day variation, regardless of whether the networks were true or inferred.

**Fig. 6.**
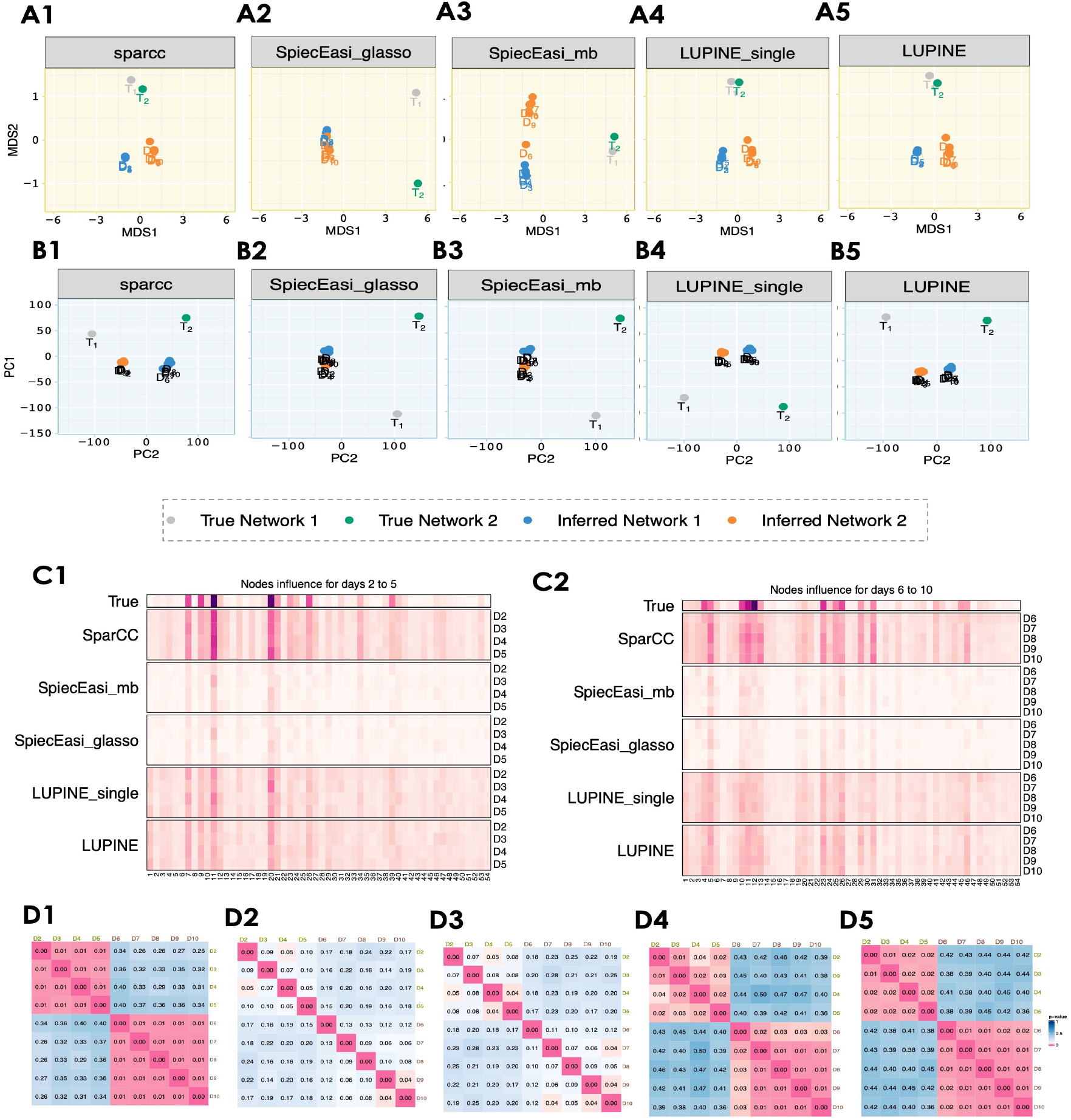
Network comparisons for simulation results n = 23. A1-A5. MDS plots of the pairwise GDD for the inferred networks from the different methods. All but SpiecEasi methods distinguish networks based on the two periods (days 2-5 and 6-10) in the first coordinate, regardless of the network being true or inferred. **B1-B5** PCA plots of the IVI scores for the inferred networks for each method. The largest node influence variation occurs across days, with a clear contrast between the two periods (days 2-5 and 6-10) for both LUPINE models and SparCC. Heatmaps of mean IVI scores for each node **C1** from day 2 to 5 and **C2** from day 6 to 10. The first row displays IVI scores for true networks from 4. All methods identified the most influential nodes. **D1-D5** Heatmaps of network correlation p-values indicate large differences between networks from days 2 to 5 from days 6 to 10 in both LUPINE models and SparCC.

We applied the same procedure used for GDDs to conduct node influence based comparisons based on IVI values. Similar to the MDS plots, SparCC (Figure 6B1) and LUPINE methods (Figure 6B4 - B5) explained the greatest between-day variation in the PCA plots. While all methods identified the most influential nodes (Figure 6C1 - C2), the two SpiecEasi methods showed less pronounced influential nodes due to their sparse network inference, compared to the other methods. Both LUPINE methods and SparCC successfully distinguished between the networks from days 2 to 5 and days 6 to 10 (Figure 6D1 - D5). As expected, the correlations were significant within the 2 to 5 day and 6 to 10 day intervals, but not between days within each interval.

#### HFHS study: LUPINE highlighted more robust longitudinal network patterns than LUPINE single

We compared LUPINE and LUPINE single on the HFHS study. This study includes a small number of time points (Figure 8A), where mice were subjected to either a HFHS or a normal diet.

Both LUPINE single and LUPINE differentiated the diet groups in the first axis of the MDS and PCA plots (Figure 7). For LUPINE single, microbial networks from the normal diet group and the HFHS diet at day 0 were clustered, indicating a similar network structure at the onset of the two different diets (Figure 7A2). However, we did not observe a close cluster of microbial networks within the normal diet networks in the PCA plot of the IVI scores (Figure 7A2), suggesting that the node influence within the normal diet group is changing over time.

**Fig. 7.**
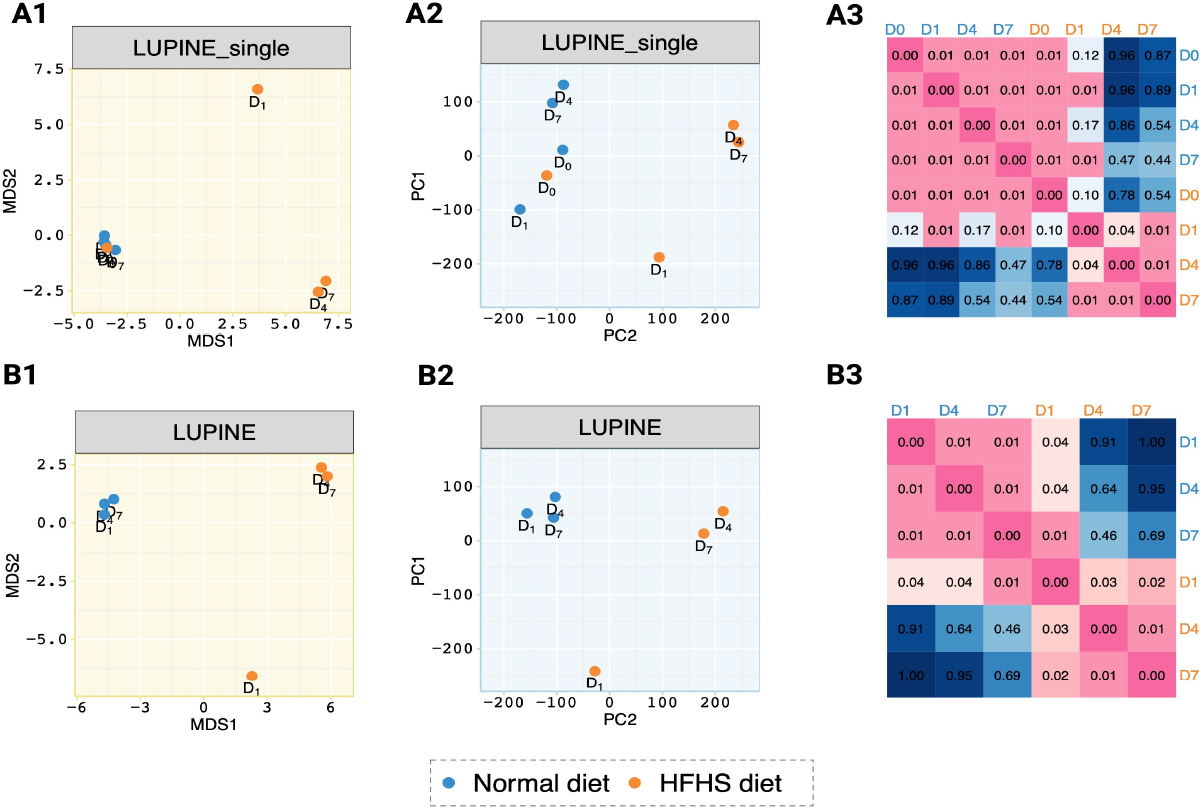
Network comparison results for LUPINE single and LUPINE. **A1-B1** MDS plot of pairwise GDD and **A2-B2** PCA plot of the IVI scores illustrating network similarities. **A** from LUPINE single scheme and **B** from LUPINE longitudinal scheme. Compared to LUPINE single, LUPINE exhibits consistent grouping patterns for both types of measures. **A3-B3** Heatmaps of network correlation p-values show that both approaches effectively identify significant differences between networks in the normal diet group and the later-stage networks in the HFHS diet group.

**Fig. 8.**
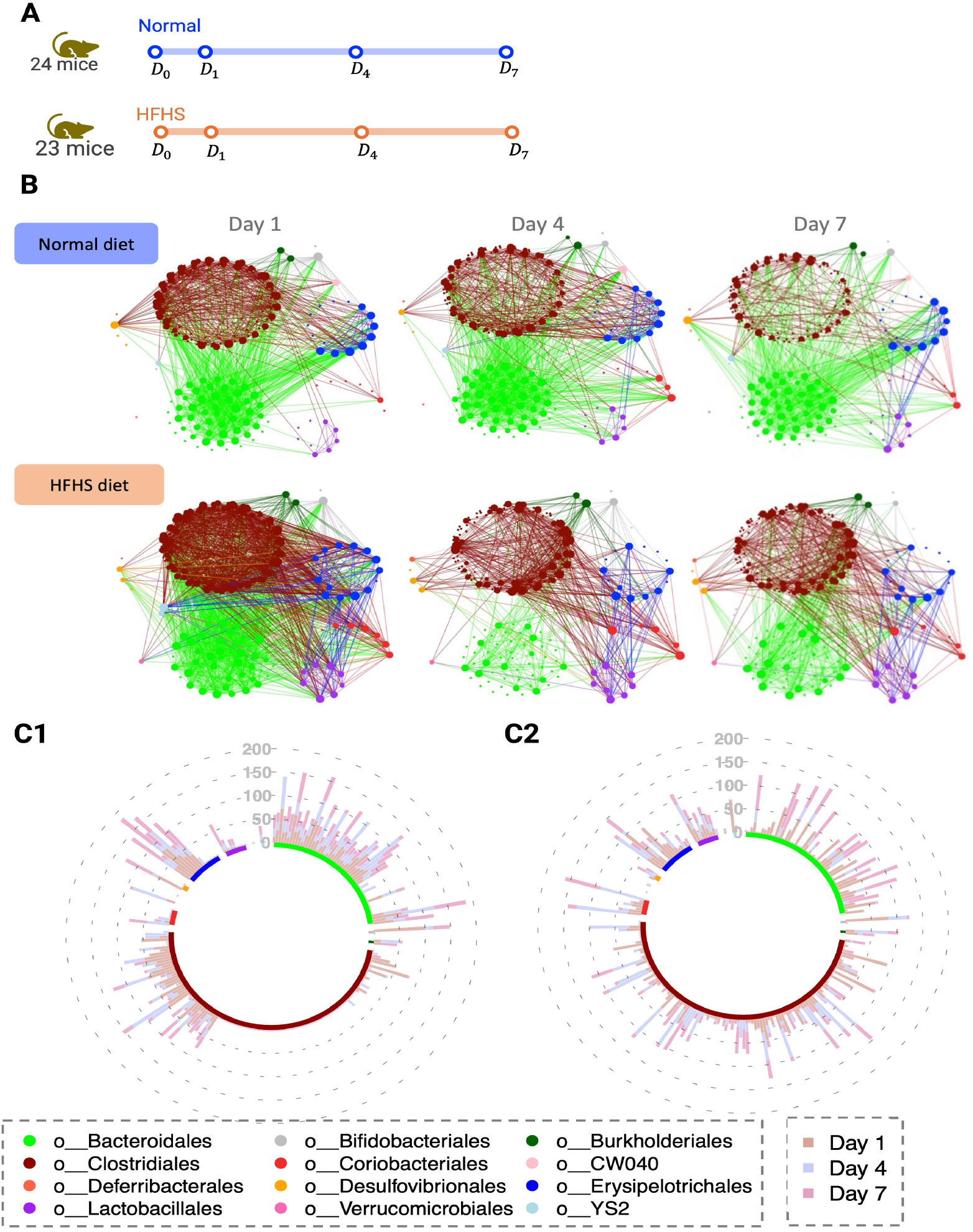
HFHS case study. **A** Graphical representation of the study timeline. **B** Inferred microbial networks over time for both normal group and HFHS diet groups. In the HFHS diet group, we observe a decrease in connections among nodes belonging to the *Bacteroidales* order over time. In the normal diet group, nodes belonging to the *Erysipelotrichales* order are more connected to those from the *Bacteroidales* order, but in the HFHS diet group, we observe that nodes belonging to the *Erysipelotrichales* order are more connected to those from the *Lactobacillales* order. **C** Circular stacked bar plot of the IVI scores for **C1** normal diet and **C2** HFHS diet groups show a reduction in IVI scores for nodes associated with the *Clostridales* order in the normal diet group compared to the HFHS diet group.

In contrast, for LUPINE, the groupings in the PCA plot (Figure 7B1) closely matched to those observed in the MDS plot (Figure 7B2), indicating consistent network results based on GDD and IVI scores. Thus, these results suggest that LUPINE is able to model robust longitudinal patterns. Note that the initial day 0 microbial network is absent from the longitudinal scheme as we require at least one prior time point for inference. However, we found that both approaches successfully identified significant correlations within each diet group across days, while correlations between diet groups across days were insignificant, particularly later in the diet (Figure 7A3-B3).

In this benchmarking section, we first demonstrated the superior model performance of both LUPINE and LUPINE single using AUC-ROC and AUC-PRC values. We then highlighted the advantages of LUPINE in comparison to our own LUPINE single approach. We showed that LUPINE led to robust longitudinal network patterns. The following sections focus on the biological interpretation of the microbial communities identified by LUPINE in four case studies.

#### HFHS study: LUPINE identified taxonomic orders that differentiate microbial networks between different diet groups in mice across four time points

We identified patterns in the microbial network plots that distinguished the two diet groups (Figure 8B). In the normal diet group, we observed denser connections among nodes within the *Bacteroidales* order, particularly on days 4 and 7, compared to the HFHS diet group. Additionally, nodes within the *Lactobacillales* order exhibited a higher number of connections in the HFHS diet group, specifically with nodes within the *Erysipelotrichales* and *Clostridales*. Similar inferences can also be made from the IVI scores in Figure 8C1 - C2: In the normal diet group, we observed decreased IVI scores across days for nodes within the *Clostridales* order, with the majority having a zero IVI value. Compared to the normal diet group, the HFHS diet group networks had a higher node influence in *Lactobacillales* order (Figure 8C2). While these findings do not imply causation, they are consistent with previous studies that reported decreased relative abundances of *Bacteroidales* and enrichment of *Lactobacillales* in mice that had been fed with a high-fat diet (Guo et al, 2017; Yin et al, 2018), suggesting a potential influence of diet on these taxonomic orders. Daniel et al (2014) also found that the proportions of *Lactobacillales* and *Erysipelotrichales* were higher in mice fed with high-fat diet. *Lactobacillus* has also been extensively studied to prevent or treat type 2 diabetes mellitus (Lee et al, 2021), indicating its potential effect on high sugar diet.

To summarise, LUPINE analysis highlighted two different microbial community networks between normal and HFHS diets, with an increase in connections in the taxa nodes belonging to *Lactobacillales* in mice that were fed with a HFHS diet.

#### VREfm study: LUPINE highlighted changes in the network structure across two abrupt interventions in a mouse study with 11 time points

In the VREfm (Figure 9A), mice underwent two interventions: antibiotic administration and VRE colonisation. Similar to the first case study, this is a much more controlled experiment compared to human studies. However, in this study the time periods are divided into three phases: naive, antibiotic, and VRE, reflecting different stages of the experiment. In contrast to the first case study, where the objective was to identify the group differences the objective here is to explore network differences between the phases.

**Fig. 9.**
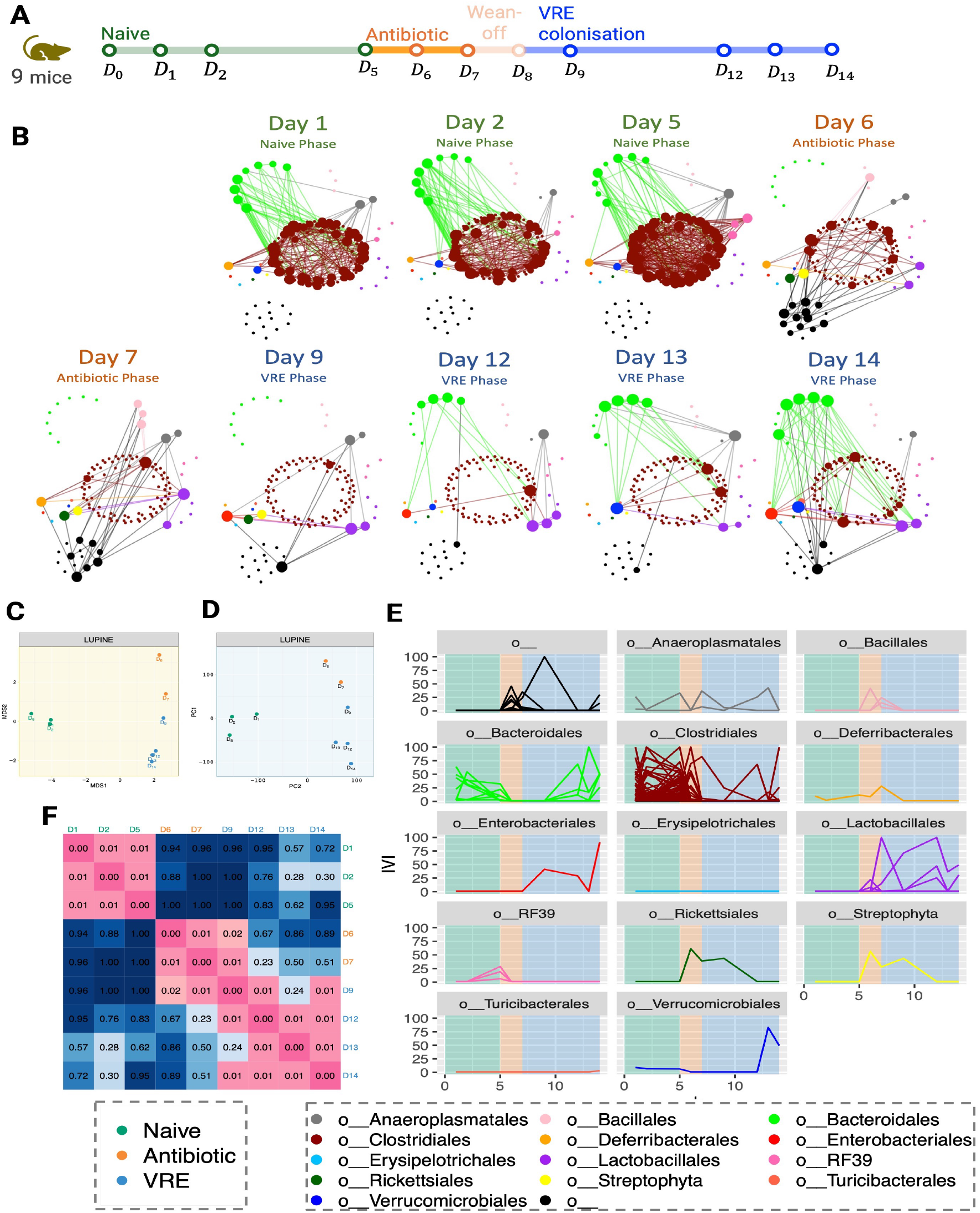
VREfm case study. **A** Graphical representation of the study timeline. **B** Inferred networks across time. After antibiotic treatment on day 6, we observe a reduction in the associations in the nodes belonging to the *Clostridales* order, which are not recovered until day 14. However, the connections among nodes of the *Bacteroidales* order appear to recover from day 12. **C** MDS plot of the pairwise GDD shows that the network structures are significantly different for each phase of the experiment. **D** PCA plot of the IVI scores exhibits a grouping pattern consistent with the MDS plot. **E** Line plots of the IVI scores for each taxa across time, grouped by their taxonomic order show a reduction of influential nodes after antibiotics treatment, indicating a less dense network structure. **F** The heatmap of Mantel p-values comparing network correlations indicates significant correlations within the naive phase, between the antibiotic phase and day 9, and across the last three days of the VRE phase.

During the antibiotic phase, the number of edges in the inferred networks decreased in *Bacteroidales* and *Clostridiales* (Figure 9B). *Bacteroidales* are an important microbe for short-chain fatty acids. Miao et al (2020) found that *Bacteroidales* almost disappear from mouse feces during ceftriaxone treatment. During the VRE phase, the number of edges increased, but this increase was not observed across all taxonomic orders. Specifically, we observed an increased number of edges in the nodes belonging to *Bacteroidales*. In contrast, and even at the end of the experiment, the nodes belonging to *Clostridiales* had very few connections compared those detecting in the naive phase. From the antibiotic phase to day 9, two nodes belonging to taxonomic order *Streptophyta* and *Rickettsiales* exhibited higher connections. This finding is in agreement with Lankelma et al (2016), who studied the disruption of the gut microbiota after the administration of broad-spectrum antibiotics and found a strong relative increase in bacteria of the genus *Streptococcus* across all subjects. Therefore, the increased connections of *Streptophyta* and *Rickettsiales* could potentially be attributed to the impact of antibiotics on the gut microbiota. From day 12, we also observed an increased number of connections in *Verrucomicrobiales*. The genome of *Akkermansia muciniphila*, a species within this taxonomic order, has been examined for its potential antibiotic resistance-associated genes (Van Passel et al, 2011).

The MDS plot of the pairwise GDD revealed a clear separation between the naive phase, antibiotic phase, and VRE phase networks (Figure 9C). Within the VRE phase, we observed a temporal progression, indicating a unique network structure on day 9, which marked the initiation of VREfm colonisation. The PCA plot of the IVI scores (Figure 9D) showed a similar grouping structure to the MDS plot. The similarity between the two plots indicated a consensus in the network grouping, regardless of the two metrics employed (as we showed previously in the HFHS study). The naive phase networks IVI scores were well separated from the other phases, accounting for 35% of total variation of the scores. After the naive phase, we observed a decrease in the influential nodes for the majority of taxa. However, taxa belonging to the *Enterobacterales* and *Lactobacillales* orders became more influential following the antibiotic phase (Figure 9E). In Kodikara et al (2022), we also found that these taxa belonging to families *Enterobacteriaceae* and *Enterococcaceae* showed a group difference between the naive and the VRE phases. We also noted a decrease in the IVI scores for *Clostridiales* nodes following the naive phase, a decrease that persisted until the end of the experiment. This taxonomic order was a key taxa group that has been studied for its ability to restrict gut colonisation by *Enterobacteriaceae* pathogens in mice, as shown by Byndloss et al (2017), Kim et al (2017), and Djukovic et al (2022). When comparing networks, we observed strong correlations within the naive phase, between the antibiotic phase and day 9, and during the later days of the VRE phase (Figure 9F). Additionally, we observed a weak correlation between the networks across phases.

To summarise, LUPINE highlighted a reduction in the number of edges after antibiotic treatment, affecting nodes belonging to the *Clostridiales* order. Interestingly, the reduction in *Clostridiales* node connections persisted even after the antibiotic phase, suggesting that the VRE colonisation had an impact on the recovery of these connections. However, further studies are needed to assess whether these connections would be recovered without VRE intervention.

#### Diet study: LUPINE detected diet-specific and diet-stable taxonomic groups in a case-control human study spanning accross 15 time points

The Diet case-control study differs from the previous ones as it involves humans and numerous time points (Fig 10A) and a small number of participants. This results in expected higher variability than mouse studies.

**Fig. 10.**
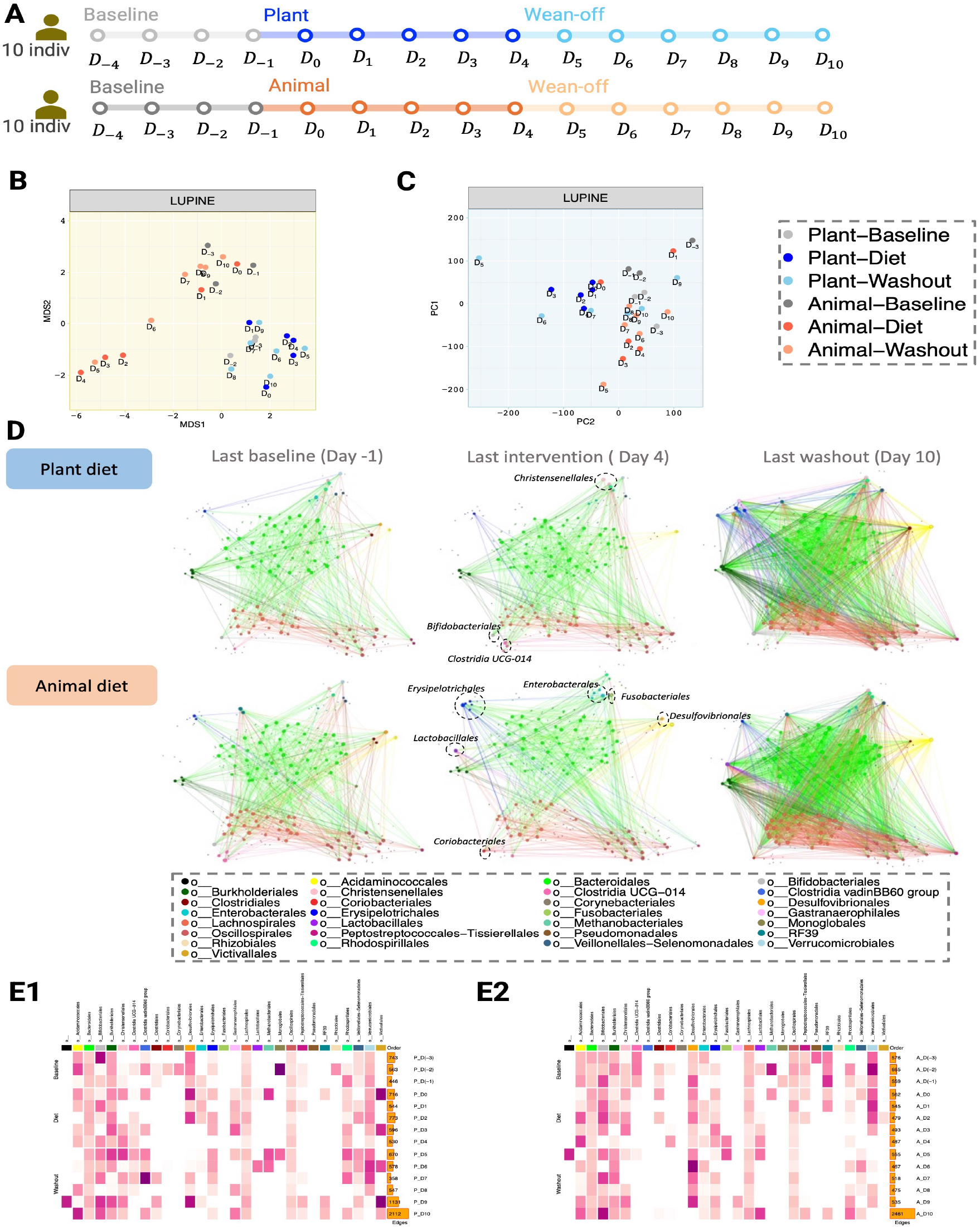
Diet case study. **A** Graphical representation of the study timeline. **B** MDS plot of pairwise GDD shows that distinct patterns emerge based on diet groups (tighter cluster for plant-based networks compared to animal-based networks). **C** PCA plot of the IVI scores show that inferred networks at baseline are similar regardless of diet, indicating similar node wise importance. **D** Inferred networks across two diet groups and three time points (last day of baseline, intervention and washout phase). On day 4, the plant-based network shows increased connections in nodes associated with *Christensenellales, Christensenellales*, and *Clostridia UCG 014* compared to the animal-based network, which exhibits increased connections in nodes related to *Erysipelotrichales, Lactobacillales, Coriobacteriales, Enterobacterales, Fusobacteriales* and *Desulfovibrionales*. **E** Heatmap of the average IVI score for each taxonomic order for **E1** plant based and **E2** animal based diet groups, with darker pink representing high average IVI value. Nodes from *Bacteroidetes, Lachnospirales* and *Oscilospirales* consistently exhibit a non-zero IVI score, indicating their stable influence, unaffected by diet or daily variations.

The plant-based diet group differed from the animal-based diet group, with the former showing a tighter network grouping, indicating more significant changes in the animal diet intervention (Figure 10B). In the animal diet group, two clusters were observed. The first included networks from days 2 to 4, and the second from days -3 to 1 and 7 to 10, with the network on day 6 distinct from both clusters. This suggests that the network structure returned to its pre-intervention form two days into the washout period. A high variability was also noted across the animal diet group networks in the PCA plot (Figure 10C). However, the PCA plot revealed a less distinct separation between the two network groupings within the animal diet group compared to the MDS plot. This indicates a larger difference in the inferred networks, due to information flow or structure, compared to the IVI score variation. Furthermore, we noticed a close alignment in the baseline inferred networks across both groups. This implies a similar degree of node influence during the baseline phase for both groups.

We examined in more detail the six inferred networks from the two diet groups at three different days, each reflecting the final day of each phase: baseline, intervention, and washout (Figure 10D). We highlighted few taxa nodes in the networks at day 4 that differed between the two diet groups. On day 4, we observed the nodes belonging to *Clostridia UCG 014* had a comparatively higher degree in the plant based diet group than in the animal based diet group. *Clostridia UCG 014* is a bacterial taxon belonging to the class *Clostridia* in the phylum *Firmicutes*. While Pessoa et al (2023) found that *Clostridia UCG 014* was reduced in the high-fat diet, Li et al (2021) associated the decrease in *Clostridia UCG 014* with obesity. Additionally, He et al (2022a) found a positive correlation between *Clostridia UCG 014* and blood glucose levels, and Jiang et al (2023) inferred *Clostridia UCG 014* as one of the taxa potentially linked to cholesterol regulation. He et al (2022b) found *Clostridia UCG 014* to be positively correlated with propionate in mice fecal samples. Propionate is a short-chain fatty acid that can be produced by gut bacteria through fermentation of dietary fiber. Additionally, *Clostridia UCG 014* was also found to be co-exist with other fiber responsive bacteria such as *Ruminococcus* (Ravelo et al, 2023). Nodes belonging to *Lactobacillales* had comparatively higher degree in the animal based diet group compared to the plant based diet group. The original study also found that the abundances of *Lactococcus lactis* and *Pediococcus acidilactici*, which belong to the taxonomy order *Lactobacillales*, significantly increased in the animal-based diet. This is in agreement with the review from Lee et al (2023) reporting an increased level of *Lactobacillus*, a member of the *Lactobacillales* order, resulting from the consumption of white meat protein (such as chicken and fish) and dairy products (like milk, cheese, yogurt, and kefir). By comparing these taxa with the average IVI scores across the intervention period, we observed some agreement between a high average IVI score (Figure 10E1 - E2) and a high number of connections in the networks in Figure 10D. For instance, *Clostridia UCG 014* had a non zero IVI average in the plant based diet group from day 2, whereas the IVI score in the animal based diet group was zero specifically during the intervention period (day 0 to day 4). We also observed that the nodes belonging to taxa order *Lactobacillales* had a non zero IVI average in the animal based diet group from day 2, whereas the IVI score in the plant based diet group was zero specifically during days 1 to 5. In the heatmaps, the bar plots revealed a high number of edges on the last day (i.e., day 10) compared to all other days in both groups. No clear reasoning can be given for this observation as there was no change at this particular time point. However, this network did not stand out in our network comparison, indicating that even though the number of connections increased, the network structure and the influential nodes on day 10 were similar to those on day 9. A densely connected network at the last time point was also visible in the network plots in Figure 10D. However, unlike the previous two case studies we did not identify diet specific correlations occurring in this particular case study. (Figure S3.3)

To summarise, LUPINE applied to a case-control human study with intervention identified several taxonomic associations that were specific to plant based diet group or animal based diet group. We also identified few taxonomic orders such as *Bacteroidetes, Lachnospirales* and *Oscilospirales* that were influential regardless of time, or diet intervention. This suggests that these taxa may exhibit stability in the face of dietary changes. In fact, *Oscillospira* is currently being explored as a next-generation probiotic due to its potential health benefits (Yang et al, 2021).

#### Pre-diabetic study: LUPINE identified key taxonomic groups studied as potential therapeutic targets for diabetic patients in a large human metagenomic study conducted at two time points

The pre-diabetic case study includes metagenomics data from a larger number of individuals in each group, across two time points (Figure 11A), with pre-diabetic individuals following either a MED diet or a PPT diet.

**Fig. 11.**
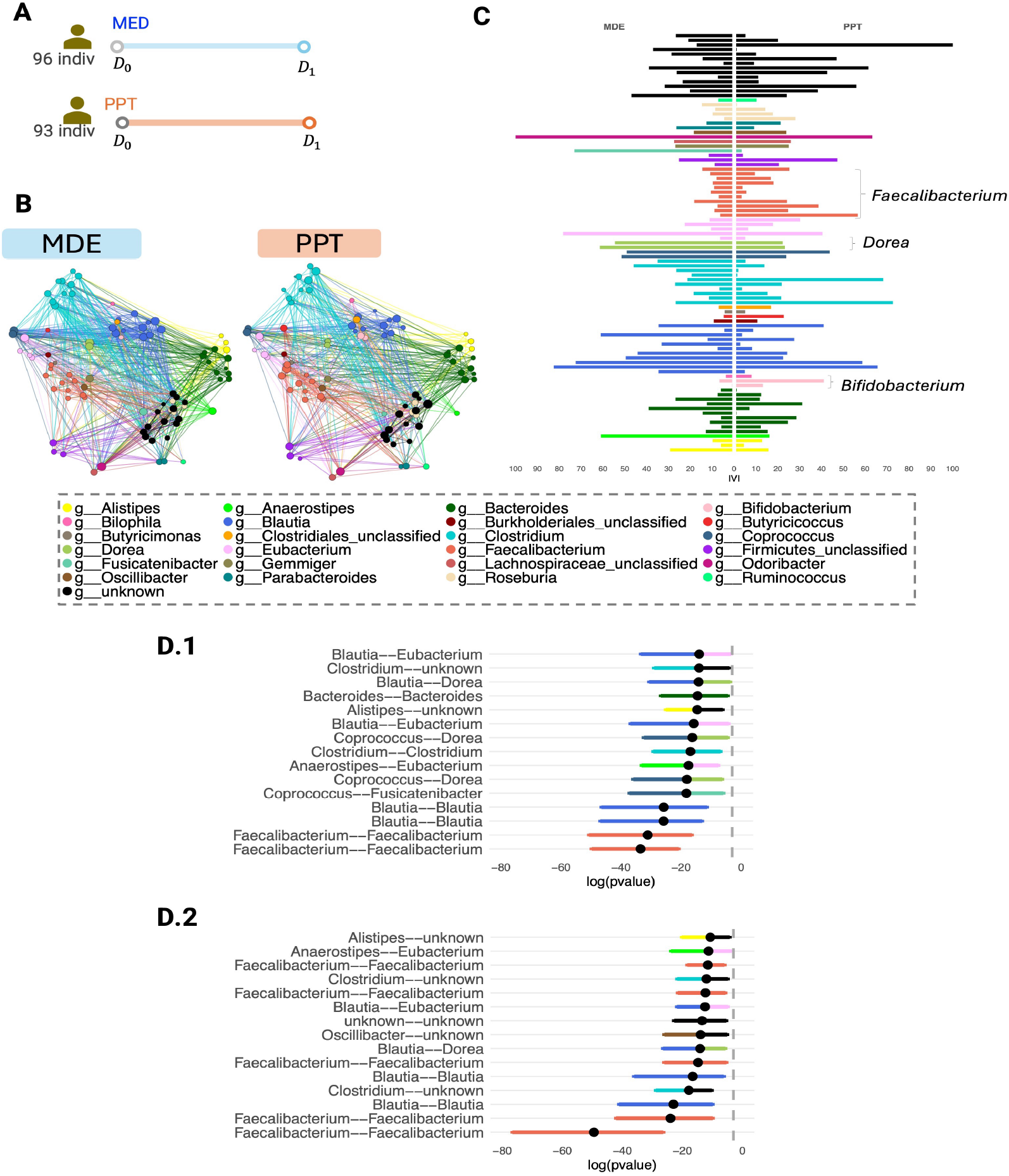
Pre-diabetic case study. **A** Graphical representation of the study timeline. **B** Inferred networks post-intervention for MDE and PPT diets show similar network structures across both diets. **C** Bar plot of IVI scores for each species-level genome bin highlights subtle differences in some genus groups. Notably, nodes belonging to*Bifidobacterium* and *Faecalibacterium* exhibit higher IVI scores in the PPT diet compared to the MDE diet. Confidence interval plots for log transformed p-values from correlation tests: **D1** MDE diet **D2** PPT diet, show a higher number of connections between *Faecalibacterium* pairs in the PPT diet compared to the MDE diet. The dashed grey vertical line represents a p-value of 0.05.

Compared to other case studies, the networks inferred through LUPINE displayed a similar network structure in the pre-diabetic case study (Figure 11B). This was further confirmed by a low p-value in the Mantel test, indicating a significant correlation between the network adjacency matrices. However, we identified differences in the two networks based on IVI scores (Figure 11C). Specifically, several genera varied between the two diet groups. For example, nodes belonging to the genus *Faecalibacterium* had higher IVI scores in the PPT group compared to the MED group. Previous research by Zhang et al (2013) showed that butyrate-producing bacteria, such as *Faecalibacterium prausnitzii*, were reduced in individuals with pre-diabetes compared to those with normal glucose tolerance. Several other studies have also reported a decline in butyrate-producing bacteria in pre-diabetic or diabetic groups (Bhute et al, 2017; Wu et al, 2020). Butyrate has been shown to down-regulate inflammation and increases mucosal barrier integrity (Chen and Vitetta, 2020). Palacios et al (2020) suggested that increased butyrate levels could enhance glucose management, while Kallassy et al (2023) demonstrated that *Faecalibacterium prausnitzii* lowered fasting blood glucose, improved glucose tolerance, and reduced HbA_1c_ levels in pre-diabetic and diabetic mice. Thus, a higher number of connections for nodes belonging to *Faecalibacterium* may indicate positive effects of the PPT diet on pre-diabetic patients.

Similarly, nodes belonging to *Bifidobacterium* also had higher IVI scores in the PPT group than in the MED group. Chang et al (2024) found lower levels of *Bifidobacterium* in pre-diabetic patients compared to healthy individuals. *Bifidobacterium* was observed as a bacterium that is indirectly capable of promoting GLP-1 (Glucagon-like peptide-1) production (Li et al, 2022). GLP-1 are a class of medication used to treat type 2 diabetes (Hinnen, 2017). Therefore, higher IVI scores for *Bifidobacterium* nodes may further suggest beneficial effects of the PPT diet on pre-diabetic patients. In contrast, nodes from *Dorea* had lower IVI scores in the PPT group than in the MED group.

Several studies have reported higher abundance of *Dorea* in pre-diabetic individuals (Allin et al, 2018; Pinna et al, 2021; Gravdal et al, 2023). Additionally, Gravdal et al (2023) identified *Dorea* as one of the top 10 bacteria differentiating healthy individuals from diabetic patients (type 2 or pre-diabetic). Positive correlations were also found between *Dorea* abundance and fasting plasma glucose, C-peptide, BMI, and waist circumference (Allin et al, 2018). Hence, lower IVI scores for *Dorea* nodes may indicate a healthier profile.

We also observed differences between the two diet groups based on the bootstrap confidence intervals of the p-values from correlation tests between pairs of taxa (Figure 11D1-D2). In the PPT diet group, most of the 15 lowest p-values indicating significant associations were between nodes belonging to *Faecalibacterium. Faecalibacterium prausnitzii*, a species from this genus, has been identified as a potential target for diabetes prevention or treatment strategies, such as prebiotics or probiotics, due to its role in producing butyrate, which enhances insulin homeostasis (Cui et al, 2022). Therefore, increased connections involving *Faecalibacterium* could reflect the positive effects of the PPT diet. In the MDE diet group, nodes belonging to *Dorea* were associated with *Coprococcus* nodes twice and with a *Blautia* node once. In the PPT diet group, *Dorea* was only associated with a *Blautia* node in the top 15 associations. Takeuchi et al (2023) showed that organisms from the genera *Blautia, Dorea*, and *Coprococcus*, as listed in the Kyoto Encyclopedia of Genes and Genomes, exhibited the three highest positive correlations with fecal carbohydrates.

To summarise, we applied LUPINE to a large case-control metagenomic human study and showed that both MDE and PPT resulted in similar networks. However, downstream analysis highlighted several taxonomic genera that were more prominent in the PPT diet, including *Faecalibacterium. Faecalibacterium prausnitzii*, a species within this genus, has been suggested as a therapeutic option for Type 2 diabetes due to its potential to enhance insulin sensitivity, improve lipid metabolism, and reduce inflammation (Xuan et al, 2023).

## 4 Discussion

We developed LUPINE to detect the stability of taxa associations within a microbial community over time, and the responses of these microbial communities to external disturbances such as dietary changes and medication. To the best of our knowledge, LUPINE is the first sequential microbial network inference approach for a longitudinal setting.

In LUPINE, we combined the concepts of low-dimensional approximation with partial correlation to infer networks across time points. We then used GDD and IVI metrics to identify any abrupt network changes across time, groups, and key taxa nodes in each of the networks. Additionally, we tested the significance of correlations between network adjacency matrices using the Mantel test. Note that a requirement for LUPINE is that samples match across time points, as LUPINE assumes correlation structure across time and individuals. In the case where different individuals are sampled across time, a better approach for network inference is our variant LUPINE single applied to each time point.

In our simulation study, we demonstrated that LUPINE and LUPINE single improved model performance and were computationally more efficient compared to existing network modelling approaches SparCC and SpiecEasi that were designed for single time point analyses. We also demonstrated the applicability of LUPINE in case studies with different study designs. The two controlled mouse studies were either case-control or intervention studies while the third and fourth studies were more complex human study incorporating elements of both a case-control and an intervention design in either long or short time courses. In all case studies, our LUPINE analyses extracted meaningful biological insights, including clustering patterns in the inferred networks that were coherent within each study design.

While developing LUPINE, we had to consider several aspects related to the characteristics of microbiome data. First, taxa filtering is an important data processing step in this kind of analysis. We only retained taxa with a mean relative abundance exceeding 0.1% for any group at any time point to obtain a consistent set of taxa being examined across various time points and groups. Further, to avoid identifying connections between taxa with low abundance at certain time points in specific groups, we then only examined taxa with a mean relative abundance greater than 0.1% within each group and time point in the inferred networks. Second, most experimental microbiome studies are typically low sample size, which are likely to result in false positive associations. We used a correlation test that is appropriate for small sample size when calculating test statistics. We chose to identify significant associations based on p-values with an arbitrary cutoff of 0.05, rather than a resampling based model selection approach (used in SpiecEasi, Kurtz et al, 2015) or a bootstrap-based p-value calculation approach (used in SparCC, Friedman and Alm, 2012) that are computationally expensive. Adjustments for false discovery on p-values could also be considered to obtain sparser networks. However, our experience has shown that these adjustments may lead to empty networks in some situations.

We have identified several potential extensions of LUPINE that could enrich longitudinal analysis of microbiome data. Firstly, when considering studies with several groups (e.g. HFHS and Diet studies), LUPINE must be applied to each group separately if we assume that the true networks within these groups differ from one another. A potential extension of LUPINE could include all groups together to infer a common network across all groups. Secondly, LUPINE main focus is to identify associations between taxa at specific time points, while taking into account information from previous time points. Therefore, we do not model time gaps between points. Future developments could include approaches that weight closer time points more heavily than distant ones. While we have only analysed one metagenomic study, which focused on taxonomy profiles based on species abundance, a future direction could be to extend this approach to include functional profiles derived from gene abundance information. An interesting extension of LUPINE to fully harness these complex data would be to establish connections between these two layers, akin to multilayer networks at each time point. This advancement could significantly broaden our understanding of microbiome dynamics, opening up new possibilities for research and discovery.

## Declarations

- Funding: National Health and Medical Research Council (NHMRC) investigator grant (GNT2025648)
- Conflict of interest/Competing interests: The authors declare they have no competing interests
- Ethics approval and consent to participate: Not applicable
- Consent for publication: Not applicable
- Data availability: All the data used in the manuscript are publicly available on Github https://github.com/SarithaKodikara/LUPINE_manuscript
- Materials availability: All materials used in the manuscript are available on Github https://github.com/SarithaKodikara/LUPINE_manuscript
- Code availability: LUPINE is available as an R package on Github https://github.com/SarithaKodikara/LUPINE.
- Author contribution: Conceived and designed the study: SK KALC. Performed the analysis: SK. Wrote the paper: SK KALC.

## References

Allin KH, Tremaroli V, Caesar R, et al (2018) Aberrant intestinal microbiota in individuals with prediabetes. Diabetologia 61:810–820

Berg G, Rybakova D, Fischer D, et al (2020) Microbiome definition re-visited: old concepts and new challenges. Microbiome 8:1–22

Bhute SS, Suryavanshi MV, Joshi SM, et al (2017) Gut microbial diversity assessment of indian type-2-diabetics reveals alterations in eubacteria, archaea, and eukaryotes. Frontiers in microbiology 8:214

Bogart E, Creswell R, Gerber GK (2019) Mitre: inferring features from microbiota time-series data linked to host status. Genome biology 20:1–15

Byndloss MX, Olsan EE, Rivera-Chávez F, et al (2017) Microbiota-activated ppar-γ signaling inhibits dysbiotic enterobacteriaceae expansion. Science 357(6351):570–575

Callahan BJ, McMurdie PJ, Rosen MJ, et al (2016) Dada2: High-resolution sample inference from illumina amplicon data. Nature methods 13(7):581–583

Chang WL, Chen YE, Tseng HT, et al (2024) Gut microbiota in patients with prediabetes. Nutrients 16(8):1105

Chen J, Vitetta L (2020) The role of butyrate in attenuating pathobiont-induced hyperinflammation. Immune network 20(2)

Cui J, Ramesh G, Wu M, et al (2022) Butyrate-producing bacteria and insulin homeostasis: The microbiome and insulin longitudinal evaluation study (miles). Diabetes 71(11):2438–2446

Daniel H, Gholami AM, Berry D, et al (2014) High-fat diet alters gut microbiota physiology in mice. The ISME journal 8(2):295–308

David LA, Maurice CF, Carmody RN, et al (2014) Diet rapidly and reproducibly alters the human gut microbiome. Nature 505(7484):559–563

Dicks LM, Geldenhuys J, Mikkelsen LS, et al (2018) Our gut microbiota: a long walk to homeostasis. Beneficial microbes 9(1):3–20

Djukovic A, Garzón MJ, Canlet C, et al (2022) Lactobacillus supports clostridiales to restrict gut colonization by multidrug-resistant enterobacteriaceae. Nature Communications 13(1):5617

Erb I (2020) Partial correlations in compositional data analysis. Applied Computing and Geosciences 6:100026

Friedman J, Alm EJ (2012) Inferring correlation networks from genomic survey data. PLoS computational biology 8(9):e1002687

Gloor GB, Macklaim JM, Pawlowsky-Glahn V, et al (2017) Microbiome datasets are compositional: and this is not optional. Frontiers in microbiology 8:2224

Gravdal K, Kirste KH, Grzelak K, et al (2023) Exploring the gut microbiota in patients with pre-diabetes and treatment naïve diabetes type 2-a pilot study. BMC Endocrine Disorders 23(1):179

Guo X, Li J, Tang R, et al (2017) High fat diet alters gut microbiota and the expression of paneth cell-antimicrobial peptides preceding changes of circulating inflammatory cytokines. Mediators of inflammation 2017

Hammond DK, Gur Y, Johnson CR (2013) Graph diffusion distance: A difference measure for weighted graphs based on the graph laplacian exponential kernel. In: 2013 IEEE Global Conference on Signal and Information Processing, pp 419–422, 10.1109/GlobalSIP.2013.6736904

He G, Chen T, Huang L (2022a) Tremella fuciformis polysaccharide reduces obesity in high-fat diet-fed mice by modulation of gut microbiota. Frontiers in Microbiology 13:1073350

He Z, Ma Y, Chen X, et al (2022b) Protective effects of intestinal gallic acid in neonatal dairy calves against extended-spectrum β-lactamase producing enteroaggregative escherichia coli infection: modulating intestinal homeostasis and colitis. Frontiers in Nutrition 9:864080

Hinnen D (2017) Glucagon-like peptide 1 receptor agonists for type 2 diabetes. Diabetes spectrum 30(3):202–210

Jiang X, Zhang B, Lan F, et al (2023) Host genetics and gut microbiota jointly regulate blood biochemical indicators in chickens. Applied Microbiology and Biotechnology 107(24):7601–7620

Jolliffe IT (1982) A note on the use of principal components in regression. Journal of the Royal Statistical Society Series C: Applied Statistics 31(3):300–303

Jolliffe IT (2002) Principal component analysis for special types of data. Springer

Kallassy J, Gagnon E, Rosenberg D, et al (2023) Strains of faecalibacterium prausnitzii and its extracts reduce blood glucose levels, percent hba1c, and improve glucose tolerance without causing hypoglycemic side effects in diabetic and prediabetic mice. BMJ Open Diabetes Research and Care 11(3):e003101

Kim YG, Sakamoto K, Seo SU, et al (2017) Neonatal acquisition of clostridia species protects against colonization by bacterial pathogens. Science 356(6335):315–319

Kodikara S, Ellul S, Lê Cao KA (2022) Statistical challenges in longitudinal microbiome data analysis. Briefings in Bioinformatics 23(4):bbac273

Kurtz ZD, Müller CL, Miraldi ER, et al (2015) Sparse and compositionally robust inference of microbial ecological networks. PLoS computational biology 11(5):e1004226

Lankelma JM, Belzer C, Hoogendijk AJ, et al (2016) Antibiotic-induced gut microbiota disruption decreases tnf-α release by mononuclear cells in healthy adults. Clinical and translational gastroenterology 7(8):e186

LêCao KA, Welham ZM (2021) Multivariate data integration using R: methods and applications with the mixOmics package. CRC Press

Lee C, Lee J, Eor JY, et al (2023) Effect of consumption of animal products on the gut microbiome composition and gut health. Food Science of Animal Resources 43(5):723

Lee YS, Lee D, Park GS, et al (2021) Lactobacillus plantarum hac01 ameliorates type 2 diabetes in high-fat diet and streptozotocin-induced diabetic mice in association with modulating the gut microbiota. Food & Function 12(14):6363–6373

Li K, Epperly MW, Barreto GA, et al (2021) Longitudinal fecal microbiome study of total body irradiated mice treated with radiation mitigators identifies bacterial associations with survival. Frontiers in Cellular and Infection Microbiology 11:715396

Li Y, Wu Y, Wu L, et al (2022) The effects of probiotic administration on patients with prediabetes: a meta-analysis and systematic review. Journal of Translational Medicine 20(1):498

Lin H, Peddada SD (2020) Analysis of compositions of microbiomes with bias correction. Nature communications 11(1):3514

Lyu R, Qu Y, Divaris K, et al (2023) Methodological considerations in longitudinal analyses of microbiome data: A comprehensive review. Genes 15(1):51

Mantel N (1967) The detection of disease clustering and a generalized regression approach. Cancer research 27(2 Part 1):209–220

Matchado MS, Lauber M, Reitmeier S, et al (2021) Network analysis methods for studying microbial communities: A mini review. Computational and structural biotechnology journal 19:2687–2698

Miao Z, Cheng R, Zhang Y, et al (2020) Antibiotics can cause weight loss by impairing gut microbiota in mice and the potent benefits of lactobacilli. Bioscience, biotechnology, and biochemistry 84(2):411–420

Mu A, Carter GP, Li L, et al (2020) Microbe-metabolite associations linked to the rebounding murine gut microbiome postcolonization with vancomycin-resistant enterococcus faecium. Msystems 5(4):10–1128

Palacios T, Vitetta L, Coulson S, et al (2020) Targeting the intestinal microbiota to prevent type 2 diabetes and enhance the effect of metformin on glycaemia: a randomised controlled pilot study. Nutrients 12(7):2041

Pessoa J, Belew GD, Barroso C, et al (2023) The gut microbiome responds progressively to fat and/or sugar-rich diets and is differentially modified by dietary fat and sugar. Nutrients 15(9):2097

Pinna NK, Anjana RM, Saxena S, et al (2021) Trans-ethnic gut microbial signatures of prediabetic subjects from india and denmark. Genome medicine 13:1–20

Ravelo AD, Arce-Cordero JA, Lobo RR, et al (2023) Effects of partially replacing dietary corn with sugars in a dual-flow continuous culture system on the ruminal microbiome. Translational Animal Science 7(1):txad011

Rothschild D, Leviatan S, Hanemann A, et al (2022) An atlas of robust microbiome associations with phenotypic traits based on large-scale cohorts from two continents. PLoS One 17(3):e0265756

Röttjers L, Faust K (2018) From hairballs to hypotheses–biological insights from microbial networks. FEMS microbiology reviews 42(6):761–780

Ruiz-Perez D, Lugo-Martinez J, Bourguignon N, et al (2021) Dynamic bayesian networks for integrating multi-omics time series microbiome data. Msystems 6(2):10–1128

Salavaty A, Ramialison M, Currie PD (2020) Integrated value of influence: an integrative method for the identification of the most influential nodes within networks. Patterns 1(5)

Sankaran K, Jeganathan P (2023) Microbiome intervention analysis with transfer functions and mirror statistics. arXiv preprint arXiv:230606364

Shoer S, Shilo S, Godneva A, et al (2023) Impact of dietary interventions on pre-diabetic oral and gut microbiome, metabolites and cytokines. Nature communications 14(1):5384

Susin A, Wang Y, Lê Cao KA, et al (2020) Variable selection in microbiome compositional data analysis. NAR Genomics and Bioinformatics 2(2):lqaa029

Takeuchi T, Kubota T, Nakanishi Y, et al (2023) Gut microbial carbohydrate metabolism contributes to insulin resistance. Nature 621(7978):389–395

Tenenhaus A, Tenenhaus M (2011) Regularized generalized canonical correlation analysis. Psychometrika 76:257–284

Torgerson WS (1958) Theory and methods of scaling. Wiley

Van Passel MW, Kant R, Zoetendal EG, et al (2011) The genome of akkermansia muciniphila, a dedicated intestinal mucin degrader, and its use in exploring intestinal metagenomes. PloS one 6(3):e16876

Waggener B, Waggener WN (1995) Pulse code modulation techniques. Springer Science & Business Media

Wold S, Sjöström M, Eriksson L (2001) Pls-regression: a basic tool of chemometrics. Chemometrics and intelligent laboratory systems 58(2):109–130

Wu H, Tremaroli V, Schmidt C, et al (2020) The gut microbiota in prediabetes and diabetes: a population-based cross-sectional study. Cell metabolism 32(3):379–390

Xuan W, Ou Y, Chen W, et al (2023) Faecalibacterium prausnitzii improves lipid metabolism disorder and insulin resistance in type 2 diabetic mice. British Journal of Biomedical Science 80:10794

Yang J, Li Y, Wen Z, et al (2021) Oscillospira-a candidate for the next-generation probiotics. Gut microbes 13(1):1987783

Yin J, Li Y, Han H, et al (2018) Melatonin reprogramming of gut microbiota improves lipid dysmetabolism in high-fat diet-fed mice. Journal of Pineal Research 65(4):e12524

Zhang X, Shen D, Fang Z, et al (2013) Human gut microbiota changes reveal the progression of glucose intolerance. PloS one 8(8):e71108

Zhang X, Pei YF, Zhang L, et al (2018) Negative binomial mixed models for analyzing longitudinal microbiome data. Frontiers in Microbiology 9:1683

Zhang X, Guo B, Yi N (2020) Zero-inflated gaussian mixed models for analyzing longitudinal microbiome data. Plos one 15(11):e0242073

