## Supplemental for "Microbial network inference for longitudinal microbiome studies with LUPINE"

### S1 Assessing the efficacy of the approximation used to improve the computation time

In this section, we evaluate the adequacy of the approximation used to improve computatoinal time using the normal diet data on day 0 from the HFHS case study ($X_0$). As a reminder, our method involves the computation of one-dimensional approximations for $p-2$ taxa. In the simplest case of a single time point, we achieved this by systematically excluding pairs of taxa from the initial dataset $X_0$, performing a principal component analysis (PCA) and extracting the first principal component. Given that there are $p \times (p-1)/2$ possible ways of selecting the excluding taxa pair, we performed PCA $p \times (p-1)/2$ on $X_0[, -(i,j)]$. $X_0[, -(i,j)]$ denote the data set $X_0$ after excluding the $i$th and $j$th taxa.

To improve computational efficiency and avoid repetitive PCA computation, we employed an approximation strategy for calculating the first principal component of $X_0[, -(i,j)]$. We first compute the loading vector for the entire dataset, $X_0$, then, approximate the first principal components for $p-2$ taxa by selectively nullifying the loading weights corresponding to the excluded taxa pair, then multiplying the resulting loading vector by $X_0$.

To compare the original principal component with its approximation we used the concordance correlation coefficient. The concordance correlation coefficient evaluates the similarity between paired data by quantifying the deviation from the concordance line (45° line through the origin) [4]. A concordance correlation value close to 1 signifies that when plotting the original principal component against its approximated counterpart, the values closely adhere to the concordance line, thus indicating a high level of agreement between the original and the approximated principal component. In Figure S1.1, we observe that all the concordance correlation values are above 0.99 between the approximated and the original principal component indicating a high level of agreement between the two. In the remainder of the main article, we therefore chose to use approximated principal component to calculate the partial correlations.

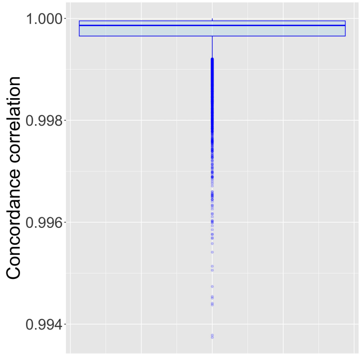


Figure S1.1: **Concordance correlation values for the normal diet data on day 0 from the HFHS case study.** The concordance correlation value is used to assess the agreement between the original and the approximated principal component for all excluded taxa pairs. All the correlation values are above 0.99 indicating a high level of agreement between the two.

### S2 Simulation Design

In this section, we describe the details of the simulation study for comparing network inference methods.

We generate data using a multivariate Poisson distribution and incorporate concepts inspired by birth and death processes [2]. At the initial time point, we use a copula-based multivariate Poisson distribution to generate 54 taxa counts for each individual. For each subsequent time point, we simulate the occurrence of new births using a multivariate Poisson process. We then update the taxa counts by adding these new births to the previous counts, which are reduced to simulate the concept of 'death'. To simulate 'death', we incorporate the First-order Integer-valued Autoregressive (INAR(1)) mechanism.

This mechanism employs a binomial thinning operator to model non-negative integer-valued time series [5]. This operator represents the decrease in taxa population between two time points. To understand the idea of a binomial thinning operator, consider a population of size $Y$ at a specific time $t$. If we observe the same population at time $t+1$, the population may decrease due to the death of some entities between times $t$ and $t+1$. Assuming these deaths are independent events and that the probability of dying between $t$ and $t+1$ is uniform, denoted by $1-\alpha$, then the resulting count of survivors at $t+1$ can be expressed as $\alpha \circ Y$ [3]. Thus, at a given time point $t$, taxa count matrix, $C_t$, is modelled as

$$C_t = \alpha \circ C_{t-1} + N_t \tag{1}$$

where $C_{t-1}$ represents the taxa count matrix at time $t-1$; $\alpha$ is the survival rate; and $N_t$ is the new birth matrix at time $t$. The matrices $C_t$, $C_{t-1}$, and $N_t$ correspond to $n$ individuals and 54 taxa.

To preserve the correlation structure, we use a correlated binomial distribution for thinning, with $\alpha = 0.5$ until time point 6 and $\alpha = 0.1$ thereafter. We assume the expected value of each taxon $i$ for individual $j$ ($\lambda_{ij}$) in the multivariate Poisson distribution remains constant over the two time periods: days 1-5 and days 6-10. We generate these $\lambda_{ij}$ values using two different Gamma distributions. For the first period (days 1-5), we generate $\lambda_{ij}$ using a Gamma distribution with its parameters estimated from the mean taxon values in the normal diet group. For the second period (days 6-10), we generate another $\lambda_{ij}$ using a different Gamma distribution with its parameters estimated from the mean taxon values in the HFHS diet group. To assess the goodness of fit between the observed mean taxon values and the fitted Gamma distribution, we generate quantile-quantile (Q-Q) plots (Figures S2.1 and S2.2). These Q-Q plots confirm the goodness of fit for the majority of taxa, as evidenced by the close alignment between the points and the theoretically expected values of the distribution.

Finally, to mimic the library size effect in our simulation, we rarefy the samples to be between 5000 and 8000.

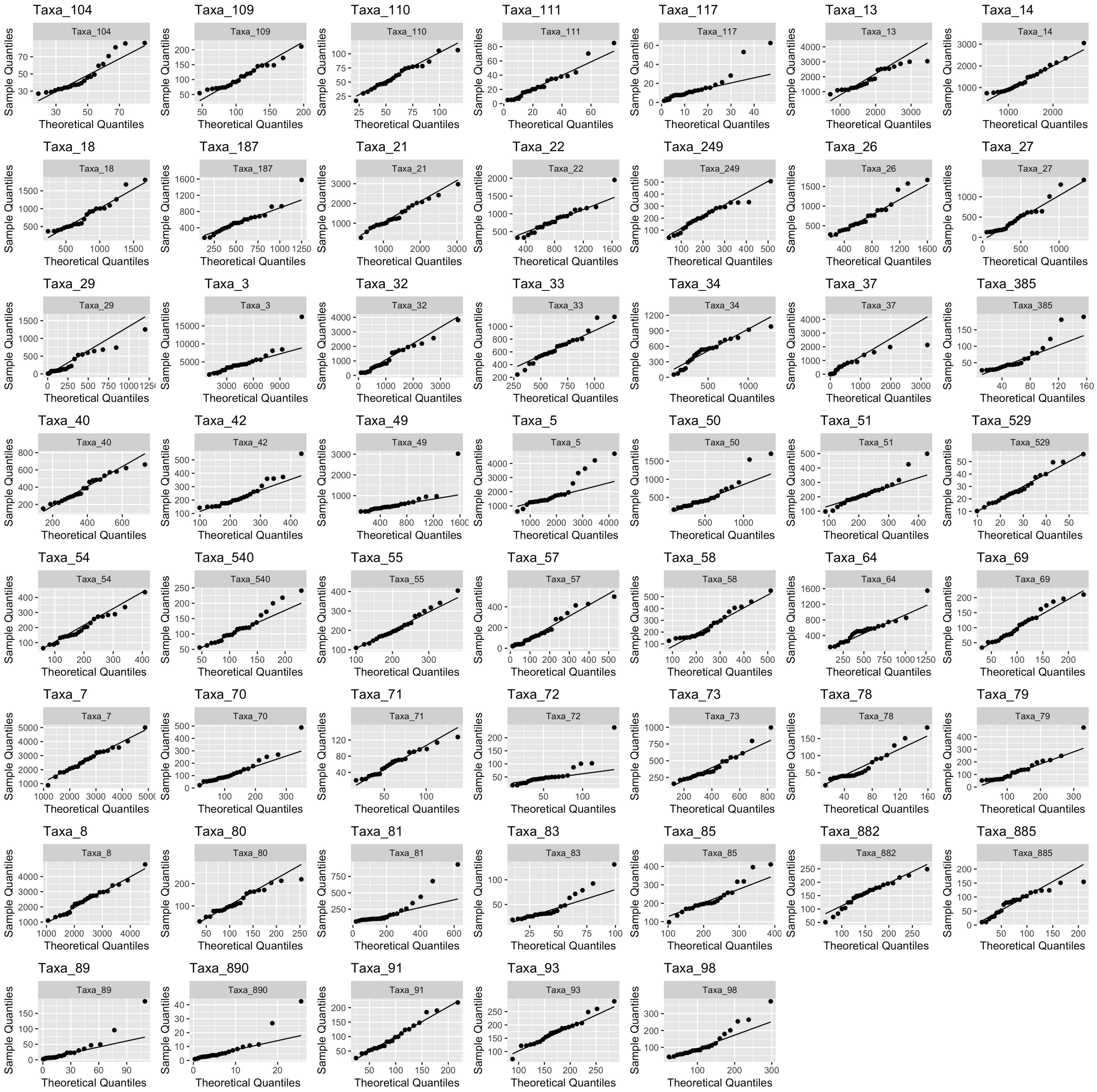


Figure S2.1: **Goodness fit of the mean taxon levels estimated from gamma distributions in the normal diet group.** For each taxon, qq plots are generated from theoretical quantiles and sample quantiles. Overall the fit of gamma distribution is satisfactory except for a few that deviate more from the theoretical quantiles (i.e. $Taxa_5$).

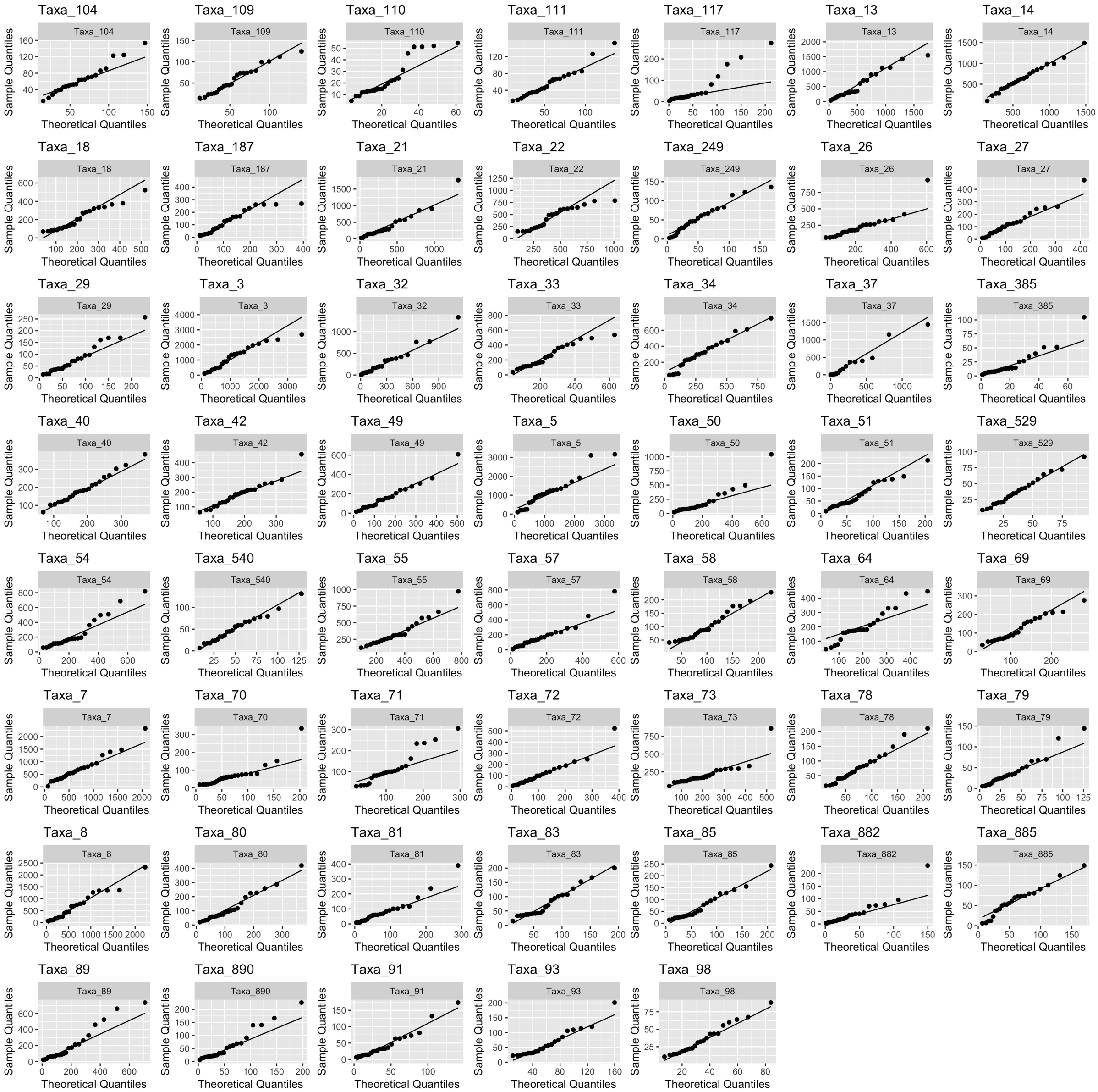


Figure S2.2: **Goodness fit of the mean taxon levels estimated from gamma distributions in the HFHS diet group.** For each taxon, qq plots are generated from theoretical quantiles and sample quantiles. Overall the fit of gamma distribution is satisfactory except for a few that deviate more from the theoretical quantiles (i.e. $Taxa_{117}$).

### S3 Supplementary Figures

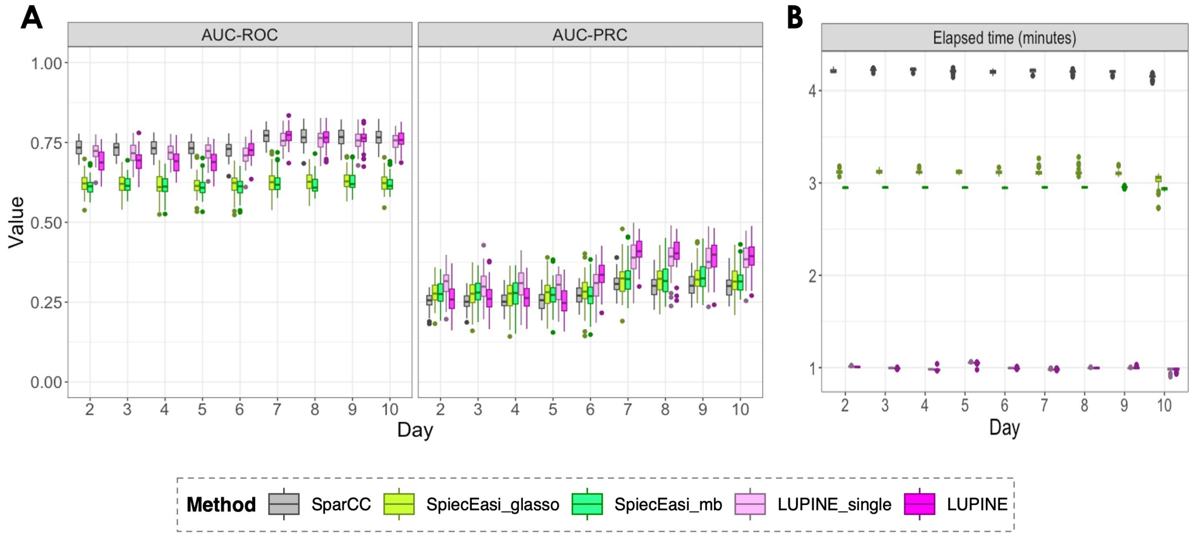


Figure S3.1: **A** Box plots of the Area Under the Receiver Operating Characteristic (AUC-ROC), Precision-Recall Curve (AUC-PRC) and **B** elapsed time values across different network inference methods for sample size 50. Each boxplot represents a distinct method, distinguished by different colours. Similar to sample size 23, two LUPINE methods and SparCC outperformed the two SpiecEasi methods based on AUC-ROC values. However, two LUPINE methods outperformed SparCC, specifically in later days. Similar to sample size 23, elapsed time was superior in LUPINE methods.

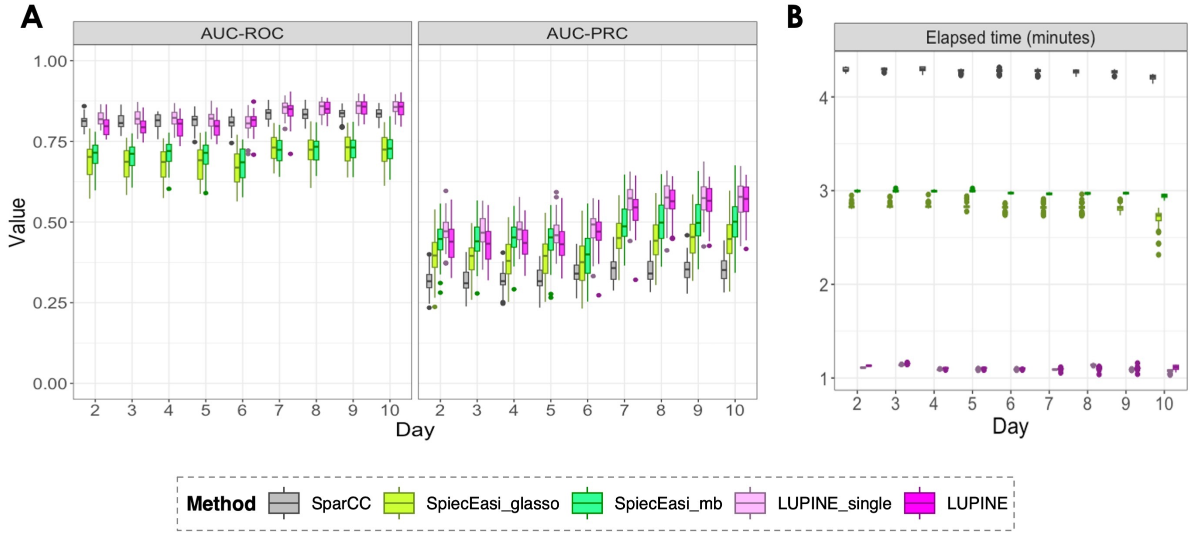


Figure S3.2: **A** Box plots of the Area Under the Receiver Operating Characteristic (AUC-ROC), Precision-Recall Curve (AUC-PRC) and **B** elapsed time values across different network inference methods for sample size 120. Each boxplot represents a distinct method, distinguished by different colours. Similar to sample size 23 and 50, two LUPINE methods and SparCC outperformed the two SpiecEasi methods based on AUC-ROC values. However, two LUPINE methods outperformed all other methods based on AUC-PRC values. Elapsed time was again superior in LUPINE methods.

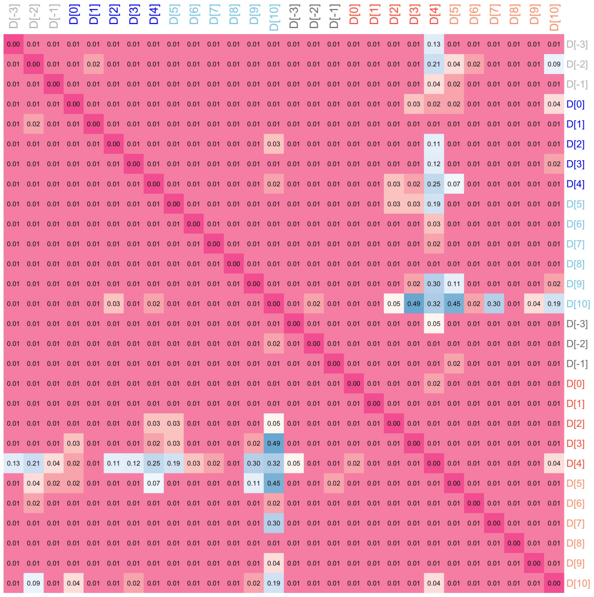


Figure S3.3: **Heatmap of Mantel p-values for diet case study** In both animal and plant diets, there is a significant correlation across networks, indicating the presence of a core network shared between the two diets.

### S4 Sensitivity Analysis

In this section, we performed a sensitivity analysis to assess the robustness of LUPINE and LUPINE_single across various conditions using simulated data. The key factors evaluated include the choice of statistical test (correlation test vs. permutation-based test), normalization techniques, number of components used in deflation, dimensionality reduction methods, edge proportions, and sample size. Our results showed that:

- Correlation-based inference were more efficient than permutation-based methods, specifically for large sample size.
- Normalization methods showed no significant impact on model performance.
- Using one component for deflation yielded comparable or superior performance to using multiple components, especially for smaller sample sizes.
- Alternative dimension reduction methods to PCA, such as robust PCA (RPCA) and independent component analysis (ICA) produced similar outcomes, with PCA showing slight advantages for larger sample sizes.
- Model performance decreased as edge proportions increased, though AUC-PRC results suggested that significant edges still captured true correlations.
- As expected, larger sample sizes enhanced model performance due to improved signal detection.

These findings reinforce the stability of LUPINE and LUPINE_single under various configurations.

#### S4.1 Permutation test vs. Correlation test

In LUPINE, we proposed using a correlation test to infer associations between taxa. On simulated data, we compared the association inferences from a permutation-based test as follows. We first calculating the partial correlation between taxa $i$ and $j$, as shown in Eq 5 (referred to as the original correlation). Next, we permuted the residuals from taxon $j$ ($e_j$) to eliminate any association between $i$ and $j$, and then calculated the correlation between $e_i$ and the permuted $e_j$. This procedure was repeated 100 times, after which we calculated the proportion of instances where the absolute value of the original correlation was less than the absolute value of the permuted correlation. A lower proportion indicated a significant association. We compared this permutation method with our correlation test results on simulated data with $n = 23$ and $n = 120$ in Section 2.5.1. As seen in Figure S4.1, correlation-based inference identified the true correlations more accurately than the permutation-based inference approach, particularly when considering the PRC values. The computational time for the permutation-based method increased exponentially with the number of nodes (i.e., taxa) as we need to perform permutations for each pair of taxa.

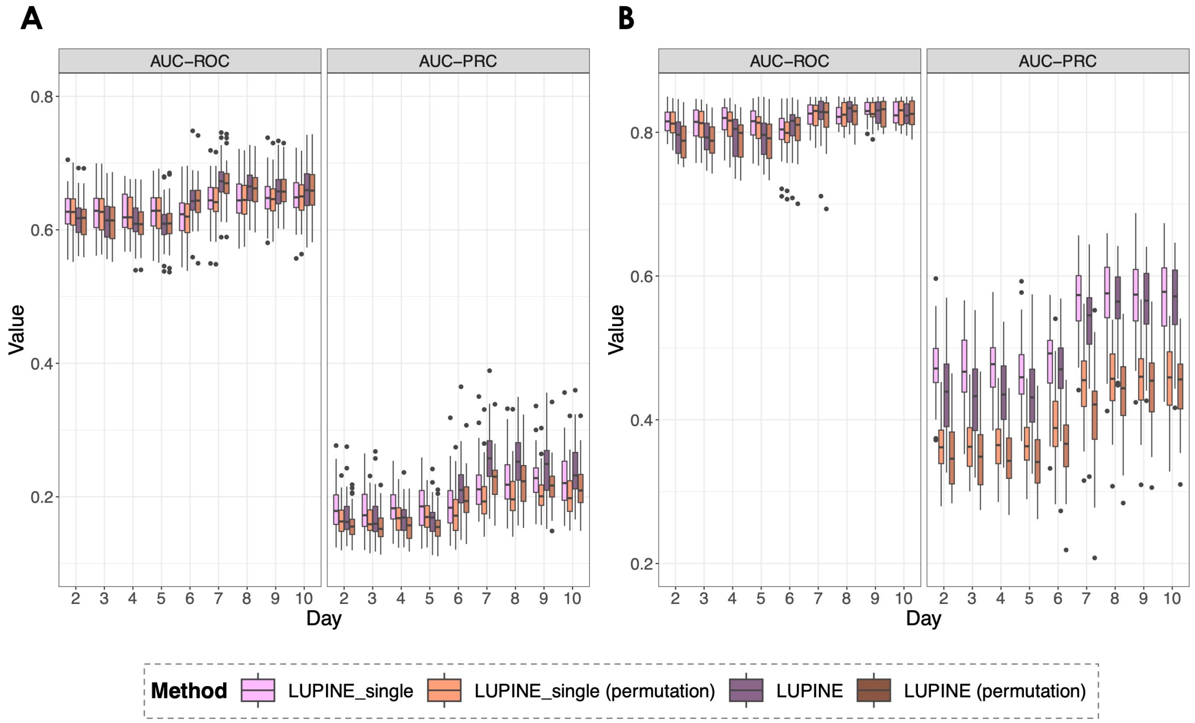


Figure S4.1: Simulated data. Box plots of the Area Under the Receiver Operating Characteristic (AUC-ROC), Precision-Recall Curve (AUC-PRC) for **A** sample size = 23 **B** sample size = 120. Each boxplot represents LUPINE_single and LUPINE evaluated with a correlation test or a permutation test, distinguished by different colours. When sample size is 23 both evaluation methods perform similarly. However, correlation based association inference outperform the permutation based inference when sample size is large.

#### S4.2 Effect of normalisation

In our original LUPINE approach, we applied PCA to the count matrix, with taxa counts centred and scaled. For the linear regressions involving taxa $i$ and $j$, we used the PCs as explanatory variables, and log-transformed the taxa counts from $i$ and $j$ to convert them to a continuous scale before using them as a response variable.

In this section, using simulated data, we evaluated the sensitivity of the results when PCA was performed on the log or clr transformed counts. Thus, in this subsection, we compared the sensitivity of LUPINE_single and LUPINE on simulated data with $n = 23$ and $n = 120$ (as described in Section 2.5.1) using the following three approaches:

1. (Default) In the linear regressions involving taxa $i$ and $j$, the explanatory variables are the log library size and the first principal component from the centred and scaled counts (excluding taxa $i$ and $j$), while the response variables are the log-transformed counts of taxa $i$ and $j$.
2. As in 1, but count data are log-transformed in PCA.
3. Explanatory variable is the first principal component from the centred and scaled clr transformed counts (excluding taxa $i$ and $j$) and the response variables are the clr transformed counts of taxa $i$ and $j$.

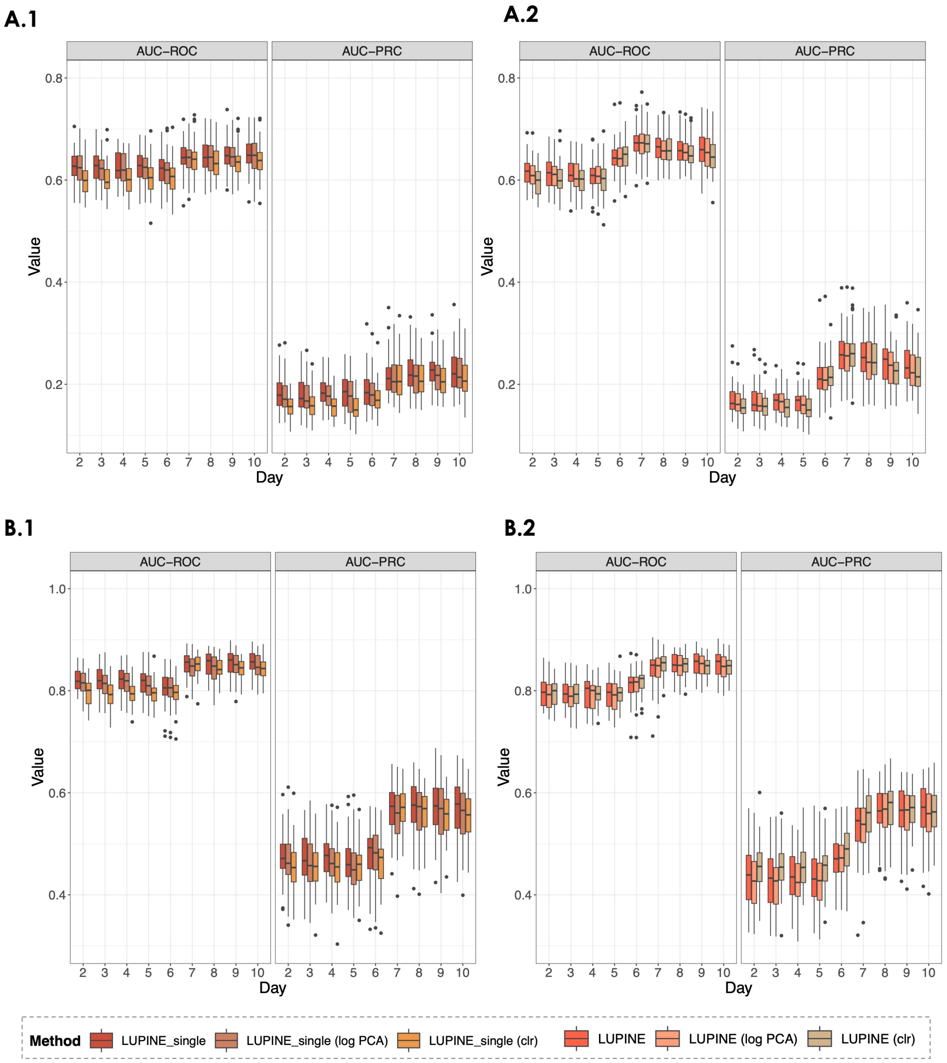


Figure S4.2: Simulated data. Box plots of the Area Under the Receiver Operating Characteristic (AUC-ROC), Precision-Recall Curve (AUC-PRC) for **A.1** sample size = 23 for different versions of LUPINE_single; **A.2** sample size = 23 for different versions of LUPINE; **B.1** sample size = 120 for different versions of LUPINE_single; **B.2** sample size = 120 for different versions of LUPINE. Each boxplot represents LUPINE_single and LUPINE evaluated with different inputs (e.g. clr tranformed vs raw counts), distinguished by different colours. For both sample sizes all three approaches produce similar results.

In the simulation, we did not observe significant differences in the three approaches considered

(Figure S4.2). Thus, in situations where count data are available, we recommend using our default version. However, users of LUPINE can choose between count or log/clr transformed data.

#### S4.3 Effect of using different number of components in the deflation

In our original approach, we used one component in the deflation to account for the largest variation in the control taxa. Including more components, however, could potentially improve model performance. To assess this, we evaluated model performance using one to three components. Our simulation results showed that using just one component produced similar or better outcomes compared to using two or three components, across two sample sizes (Figure S4.3). In particular, the one-component approach performed better when the sample size was small.

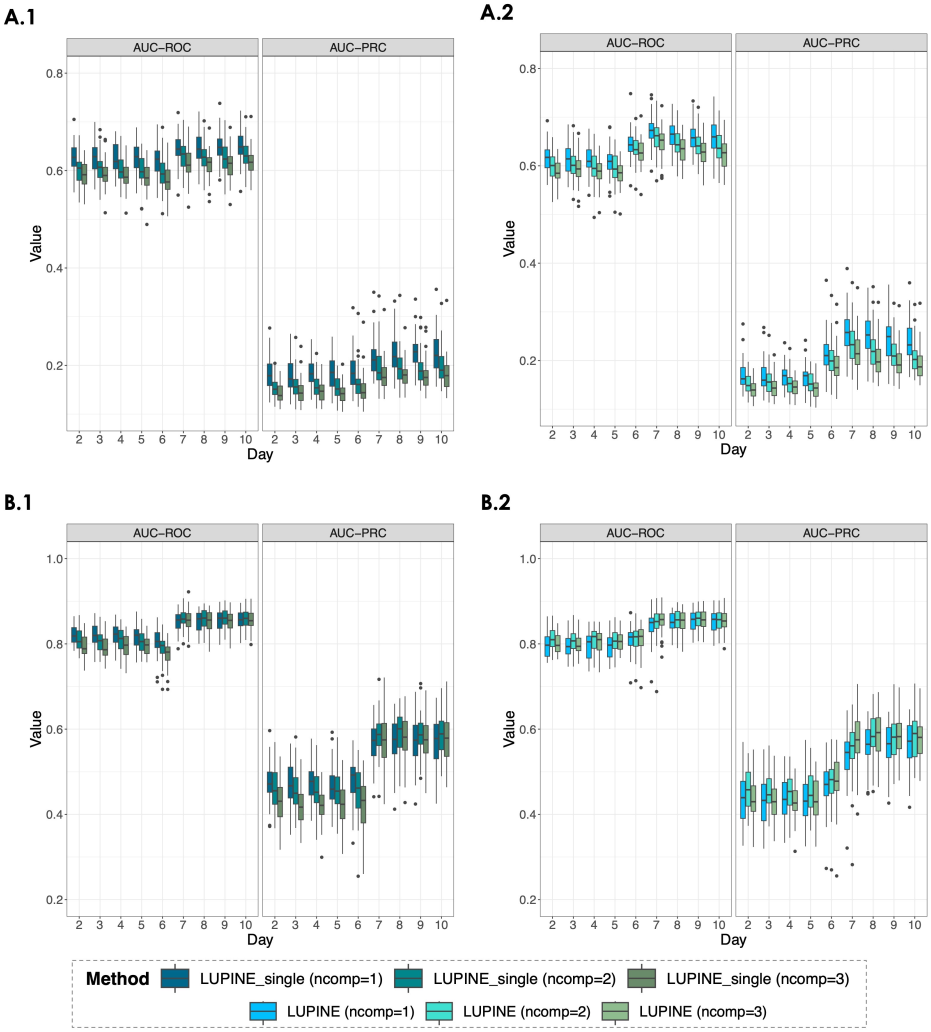


Figure S4.3: Simulated data. Box plots of the Area Under the Receiver Operating Characteristic (AUC-ROC), Precision-Recall Curve (AUC-PRC) for **A.1** sample size = 23 for LUPINE_single; **A.2** sample size = 23 for LUPINE; **B.1** sample size = 120 for LUPINE_single; **B.2** sample size = 120 for LUPINE with one to three components used in deflation. Each box plot compares the performance of LUPINE_single and LUPINE with different numbers of components used in deflation. Colors differentiate between the number of components. Results are similar for both sample sizes when using one, two, or three components.

#### S4.4 Effect of using different dimension reduction methods in LUPINE_single

In LUPINE_single, we used PCA as the dimensionality reduction method. **(author?)** [1] suggested other dimension reduction methods that might be suitable for microbiome data. We focused on RPCA, as it was available in R and could be easily integrated into our package. We also considered ICA. We obtained similar performance for PCA, ICA, and RPCA on our simulated data. However, PCA showed slightly better performance with a larger sample size ($n = 120$) (Figure S4.4). Thus, PCA is kept as the default option in the package, with alternatives for RPCA and ICA.

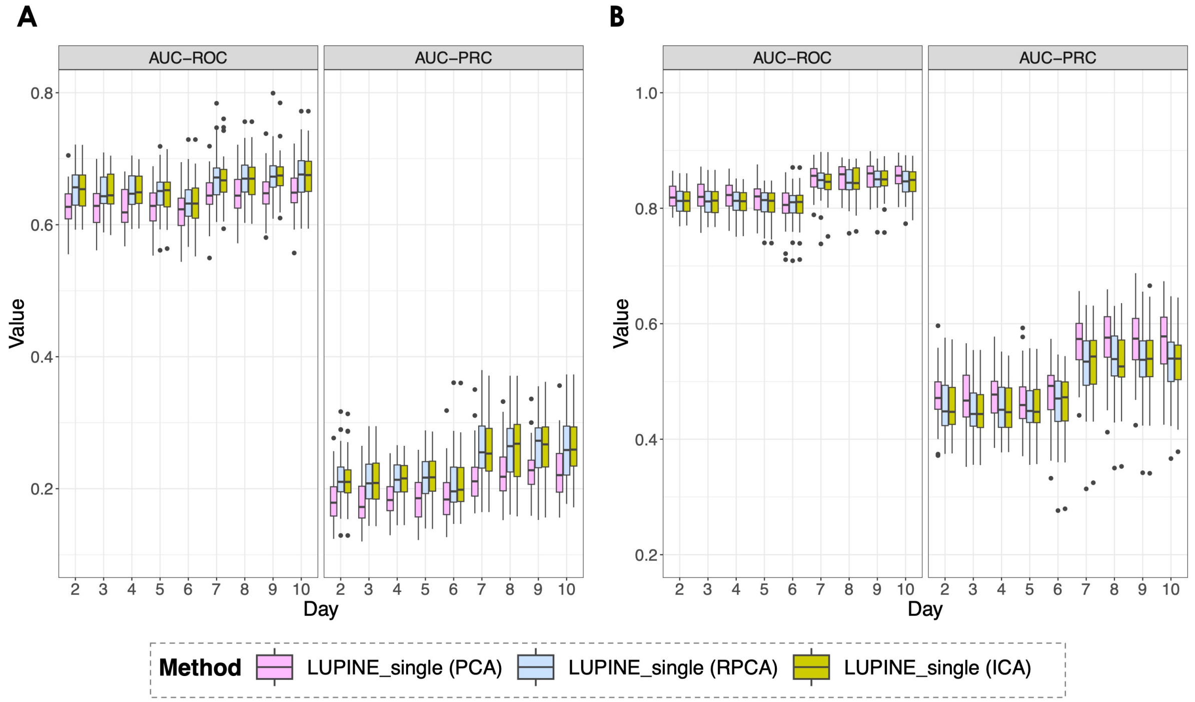


Figure S4.4: Simulated data. Box plots of the Area Under the Receiver Operating Characteristic (AUC-ROC), Precision-Recall Curve (AUC-PRC) for **A** sample size = 23 **B** sample size = 120. Each boxplot represents LUPINE_single evaluated with a PCA, RPCA, and IPCA, each distinguished by different colours. All three dimension reduction approaches produce similar results.

#### S4.5 Effect of number of edges

To evaluate the performance of the LUPINE_single and LUPINE models, we tested them using different edge proportions with a sample size of 23. Edge proportion was defined as the ratio of observed edges to the total number of possible edges. Model performance, measured by AUC-ROC, decreased as the edge proportion increased from 0.005 to 0.8 (Figure S4.5A). This decline occurred because increasing the number of edges reduced the true partial correlations. Maintaining the positive definiteness of the partial correlation matrix required this adjustment (Figure S4.5B). As partial correlations decreased, detecting significant correlations that differed from zero became more difficult, resulting in lower performance. However, performance based on AUC-PRC initially decreased and then improved, suggesting that significant edges were more likely to represent true correlations.

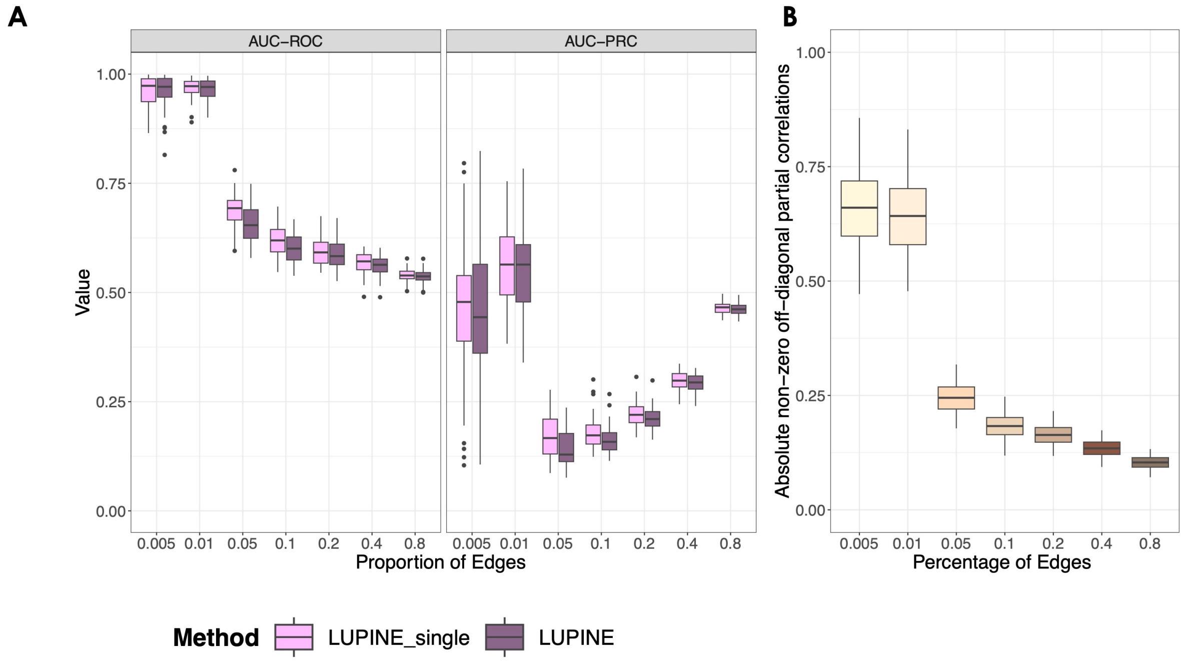


Figure S4.5: Simulated data. Box plots of the **A** Area Under the Receiver Operating Characteristic (AUC-ROC), Precision-Recall Curve (AUC-PRC) for different percentages of edges with a sample size = 23 **B** true absolute partial off-diagonal correlations. As the number of edges increases, the true correlation values decrease to maintain the positive definite property of the partial correlation matrix. Consequently, as the number of edges increases, AUC-ROC performance declines.

#### S4.6 Effect of sample size

We tested model performance across different sample sizes using an edge proportion of 0.1. The choice of a 0.1 edge proportion was inspired by the edge proportions in the simulated network shown in Figure 4A, which was inferred using SpiecEasi. Additionally, performance was generally worse at this edge proportion, as seen in the AUC-ROC and AUC-PRC values in Figure S4.5. As the sample size increased, the signal-to-noise ratio increased. Thus, as expected, the performance of both approaches improved with increasing sample size (Figure S4.6).

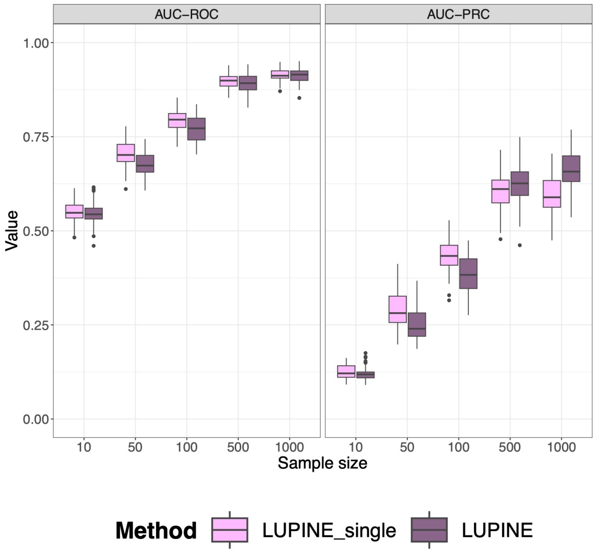


Figure S4.6: Simulated data. Box plots of the **A** Area Under the Receiver Operating Characteristic (AUC-ROC), Precision-Recall Curve (AUC-PRC) for different sample sizes with an 0.1 edge percentage. As the sample size increases, both performance measures increases.

### References


[1] George Armstrong, Gibraan Rahman, Cameron Martino, Daniel McDonald, Antonio Gonzalez, Gal Mishne, and Rob Knight. Applications and comparison of dimensionality reduction methods for microbiome data. *Frontiers in bioinformatics*, 2:821861, 2022.

[2] Willy Feller. Die grundlagen der volterraschen theorie des kampfes ums dasein in wahrscheinlichkeitstheoretischer behandlung. *Acta Biotheoretica*, 5(1):11–40, 1939.

[3] M Aghababaei Jazi and MH Alamatsaz. Two new thinning operators and their applications. *Global Journal of Pure and Applied Mathematics*, 8(1):13–28, 2012.

[4] I Lawrence and Kuei Lin. A concordance correlation coefficient to evaluate reproducibility. *Biometrics*, pages 255–268, 1989.

[5] DM Simarmata, Fevi Novkaniza, and Yekti Widyaningsih. A time series model: First-order integer-valued autoregressive (inar (1)). In *AIP Conference Proceedings*, volume 1862. AIP Publishing, 2017.
